# Dynamic chromosomal interactions and control of heterochromatin positioning by Ki-67

**DOI:** 10.1101/2021.10.20.465140

**Authors:** Tom van Schaik, Stefano G. Manzo, Athanasios E. Vouzas, Ning Qing Liu, Hans Teunissen, Elzo de Wit, David M. Gilbert, Bas van Steensel

**Affiliations:** Division of Gene Regulation and Oncode Institute, Netherlands Cancer Institute, Plesmanlaan 121, 1066 CX Amsterdam, the Netherlands; Department of Biological Science, The Florida State University, Tallahassee, 32304, FL, USA; San Diego Biomedical Research Institute, 3525 John Hopkins Court, San Diego, 92121, CA, USA; Department of Cell Biology, Erasmus University Medical Centre, Rotterdam, the Netherlands

## Abstract

Ki-67 is a chromatin-associated protein with a dynamic distribution pattern throughout the cell cycle, and is thought to be involved in chromatin organization. Lack of genomic interaction maps has hampered a detailed understanding of its roles, particularly during interphase. By pA-DamID mapping in human cell lines we found that Ki-67 associates with large genomic domains that overlap mostly with late-replicating regions. Early in interphase, when Ki-67 is present in pre-nucleolar bodies, it interacts with these domains on all chromosomes. However, later in interphase, when Ki-67 is confined to nucleoli, it shows a striking shift towards small chromosomes. Nucleolar perturbations indicate that these cell cycle dynamics correspond to nucleolar maturation during interphase, and suggest that nucleolar sequestration of Ki-67 limits its interactions with larger chromosomes. Furthermore, we demonstrate that Ki-67 does not detectably control chromatin-chromatin interactions during interphase, but it competes with the nuclear lamina for interaction with late-replicating DNA, and it controls replication timing of (peri)centromeric regions. Together, these results reveal a highly dynamic choreography of genome interactions and roles for Ki-67 in heterochromatin organization.

## INTRODUCTION

Ki-67 is a chromosomal, nuclear and nucleolar protein that is widely used as a marker for cellular proliferation (reviewed in Remnant *et al*, 2021; Scholzen & Gerdes, 2000; Sun & Kaufman, 2018). It has been implicated in chromatin biology in various stages of the cell cycle. During mitosis, Ki-67 is a key component of the peri-chromosomal layer (PCL) (Booth *et al*, 2014; Verheijen *et al*, 1989), where it acts as surfactant to prevent chromosomal intermingling (Cuylen *et al*, 2016; Takagi *et al*, 2016). Following anaphase, Ki-67 changes from a repelling into an attracting behavior to exclude cytoplasmic proteins and compact chromosomes (Cuylen-Haering *et al*, 2020). Early in interphase Ki-67 accumulates in pre-nucleolar bodies (PNBs), which are punctate structures containing rRNA precursors and various proteins (Ochs *et al*, 1985). These PNBs gradually fade away as several mature nucleoli are formed (Carron *et al*, 2012; Dundr *et al*, 2000; Savino *et al*, 2001). In these mature nucleoli Ki-67 is positioned specifically at the nucleolar rim.

Together with the nuclear lamina (NL), the nucleolus is a major hub for heterochromatin, as illustrated by both microscopy (Lima-De-Faria & Reitalu, 1963; Ohno *et al*, 1959) and genomics observations (Dillinger *et al*, 2017; Guelen *et al*, 2008; van Koningsbruggen *et al*, 2010; Vertii *et al*, 2019). Often, individual heterochromatic genomic loci are stochastically distributed between the NL and nucleoli, with variable preference for one or the other (Kind *et al*, 2013; Ragoczy *et al*, 2014; Vertii *et al.*, 2019). Additionally, disruption of one of the two structures may enhance interactions with the other (Ragoczy *et al*., 2014; Solovei *et al*, 2013), which may indicate a competitive mechanism. Interestingly, depletion of Ki-67 has been shown to lead to a loss of heterochromatin around the nucleolus (Sobecki *et al*, 2016), suggesting that it may tether heterochromatin to the nucleolus.

So far, most studies of the interplay between Ki-67 and chromatin have relied on microscopy observations (e.g., Booth *et al.*, 2014; Matheson & Kaufman, 2017; Sobecki *et al.*, 2016). While these experiments have been highly informative, it has remained unclear how exactly Ki-67 interacts with the genome throughout the cell cycle. Genome-wide interaction data would greatly enhance this understanding, and permit comparisons with other nuclear positioning data, the epigenetic landscape and functional readouts of the genome such as transcription and replication timing.

Here, we provide such data using our recently developed pA-DamID technology, which allows for simultaneous in situ visualization of protein-DNA interactions and generation of genome-wide interaction maps (van Schaik *et al*, 2020). Our results uncover remarkably dynamic interactions of Ki-67 with the genome of human cells, and provide insights into its roles in heterochromatin organization and replication timing.

## RESULTS

### pA-DamID captures genome – Ki-67 interactions

We used our recently developed pA-DamID method to profile Ki-67 interactions with the genome. pA-DamID allows us to both create maps of genome-wide protein-DNA interactions, and to visualize these interactions in situ with the ^m6^A-Tracer protein (**Fig 1A**) (Kind *et al.*, 2013; van Schaik *et al.*, 2020). Following pA-DamID with a Ki-67 antibody in hTERT-RPE, HCT116 and K562 human cell lines, we indeed observe that ^m6^A-Tracer binding (and hence the interaction of Ki-67 with the genome) is enriched at nucleoli stained by Ki-67, compared to a free Dam control (**Fig 1B, C**). ^m6^A-Tracer binding occurs mostly at the edges of nucleoli, indicating that Ki-67 preferentially contacts DNA at the nucleolar periphery. However, Ki-67 is not exclusively localized at nucleoli and may be locally enriched elsewhere, such as the nuclear periphery (**Fig 1B**, orange arrows). This overlaps with ^m6^A-Tracer staining, indicating that Ki-67 at these sites can also engage in genome interactions. Despite these individual foci, on average ^m6^A-Tracer binding is not enriched at the nuclear periphery compared to the Dam control in hTERT-RPE and HCT116 cells (**Fig S1A**). In addition, a moderate homogeneous ^m6^A-Tracer signal throughout the nucleus may be caused by low concentrations of DNA-interacting Ki-67 in the nuclear interior, but also from non-specific antibody binding.

**Fig 1.**
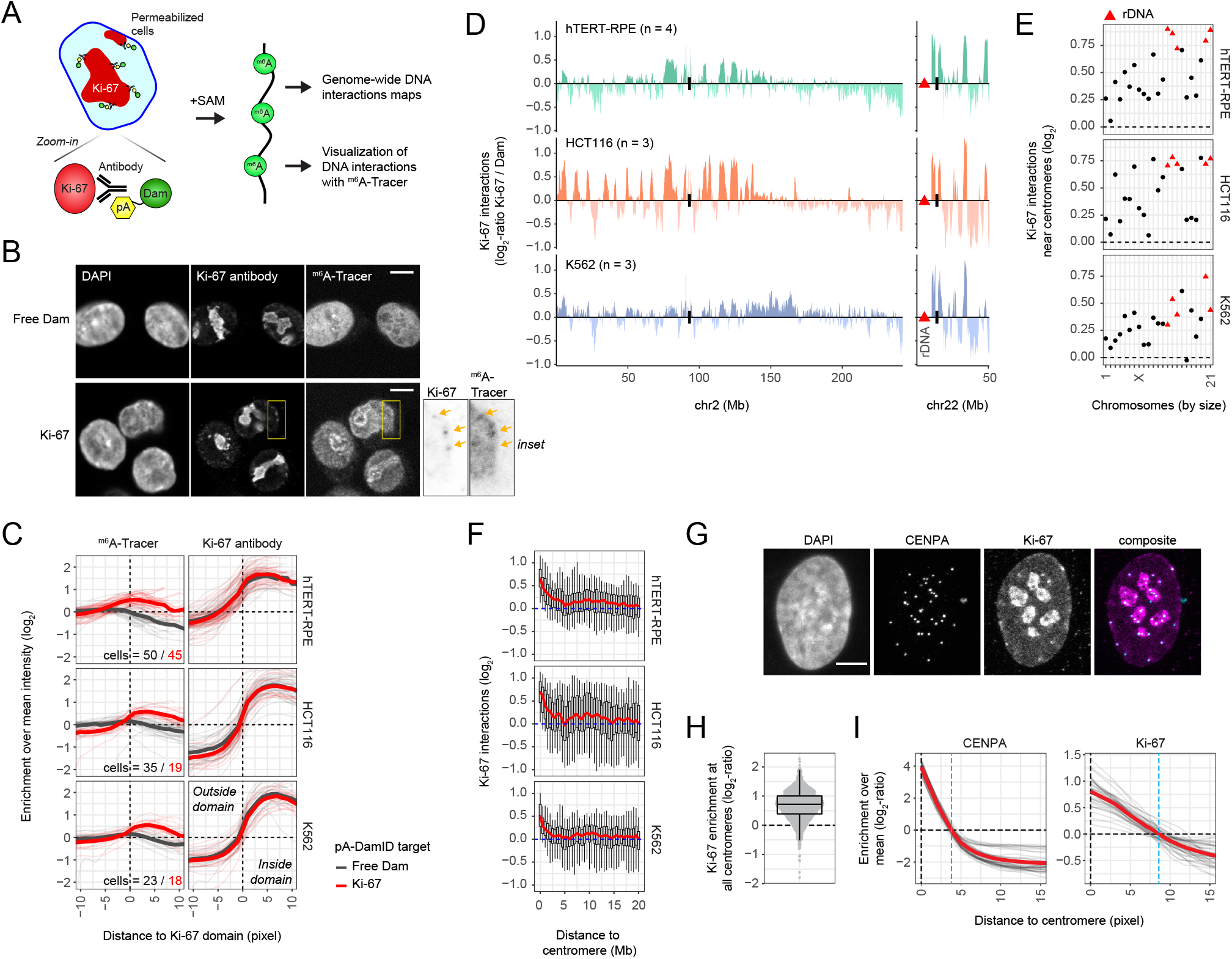
Visualization and genome-wide profiling of DNA-Ki-67 interactions using pA-DamID. **(A)** Schematic overview of pA-DamID (van Schaik *et al.*, 2020). Permeabilized cells are incubated with a primary antibody (e.g., against Ki-67), followed by a fusion of protein A and Dam (pA-Dam). After removal of unbound pA-Dam, the Dam enzyme is activated by addition of S-adenosylmethionine (SAM), resulting in local deposition of ^m6^A marks. ^m6^A-marked DNA can be processed for high-throughput sequencing, or alternatively cells can be fixed and ^m6^A marks visualized using the ^m6^A-Tracer protein. **(B)** Representative confocal microscopy sections of HCT116 cells following pA-DamID with free Dam (top panel) or Ki-67 antibody (bottom panel), labeled with ^m6^A-Tracer protein and stained for Ki-67. Scale bar: 5 μm. **(C)** Quantification of the enrichment of Ki-67 antibody and ^m6^A-Tracer signals relative to segmented Ki-67 domains (that we interpret as nucleoli) in different cell lines. For every cell, the enrichment is calculated by pixel-distances (pixel: 80 nm) relative to the mean signal of that cell, and represented as a log_2_-ratio. Negative distances are outside of Ki-67 domains, a distance of zero marks the domain boundary and positive distances are inside of Ki-67 domains. Every thin line corresponds to an individual cell and the thick line is the mean of all cells. Results are combined from three (hTERT-RPE) or one (HCT116 and K562) biological replicates. The number of analyzed cells is included in the bottom right of each ^m6^A-Tracer panel. **(D)** Comparison of Ki-67 pA-DamID profiles (log_2_-ratios over the free Dam control) across two chromosomes in hTERT-RPE, HCT116 and K562 cells. Sequenced reads are counted and normalized in 50 kb bins. Data are averages of *n* biological replicates and smoothed with a running mean across nine 50 kb bins for visualization purposes. Centromeres are highlighted by black bars. **(E)** The mean Ki-67 interaction score near centromeres is plotted for each chromosome ordered by size (within 2Mb of centromeres). rDNA-containing chromosomes are highlighted in red. **(F)** Distributions of Ki-67 interactions nearby centromeres. Boxplots are drawn for every 0.5 Mb, following the specification as in (E) with the 50^th^ percentile highlighted in red. **(G)** Representative confocal maximum projection of hTERT-RPE cells stained for CENPA and Ki-67. Cells were treated with 0.05% dimethyl sulfoxide (DMSO). Scale bar: 5 μm. **(H)** Quantification of the Ki-67 enrichment at centromeres, relative to the mean Ki-67 intensity of a cell. Every point represents one centromere. Results are combined from two biological replicates. In total, 44 cells were analyzed. **(I)** Average enrichment of CENPA and Ki-67 from centromeres. For every cell, the enrichment is calculated by pixel-distances (pixel: 80 nm) relative to the mean signal of that cell, and represented as a log_2_-ratio. Every thin line corresponds to an individual cell, and the thick line is the mean of all cells.

We then processed these ^m6^A-tagged DNA samples for high-throughput sequencing to identify the genomic regions that interact with Ki-67. We first describe results in unsynchronized cells; below we discuss the dynamics throughout the cell cycle. As described previously (van Schaik *et al.*, 2020), pA-DamID utilizes a free Dam control to normalize for DNA accessibility and amplification biases (Greil *et al*, 2006). After this Dam normalization we observed a striking domain-like pattern of Ki-67 binding to the genome (**Fig S1B**). We balanced data resolution and reproducibility by using 50 kb averaging bins that yield Pearson correlation coefficients between independent replicate experiments in the range of 0.40 - 0.80 (**Fig S1C, D**); at smaller bin sizes the data were too noisy to be informative.

To validate these interaction maps, we used an HCT116 cell line with mClover- and AID-tagged Ki-67, which allow for protein visualization and rapid protein depletion upon addition of auxin (Takagi *et al.*, 2016). Incubation of these cells with auxin for 24 hours resulted in a near-complete depletion of mClover fluorescence, but only a partial decrease in Ki-67 immunostaining signal (**Fig S1E, F**). This difference may be caused by the higher sensitivity of indirect immunofluorescence (Takagi *et al.*, 2016). In accordance with previous RNAi depletion experiments (Booth *et al.*, 2014), the residual Ki-67 signal appeared to localize in fewer and more interior positioned nucleoli (**Fig S1E**). Genome-wide mapping of Ki-67 interactions with pA-DamID resulted in a strong signal loss at the Ki-67 interaction domains upon addition of auxin (**Fig S1G, H**). We assume that the remaining signals (e.g., on chr22) result from the residual Ki-67 protein. These data verify that Ki-67 interaction domains are specific for Ki-67 and not caused by technical artifacts.

Because Ki-67 is a very large protein (~350 kDa), we reasoned that the location of the antibody epitope could affect the observed interaction patterns. Ki-67 contains protein binding domains and a DNA binding domain, positioned at the N and C-terminus, respectively (reviewed in Sun & Kaufman, 2018). The initial antibody we used was generated against a peptide sequence roughly in the middle of the protein (~1150/3256 amino acids), so we chose two additional antibodies to target each protein end. As before, immunostaining following auxin-mediated Ki-67 depletion confirms antibody specificity to Ki-67 (**Fig S2A**). Following pA-DamID, only the C-terminus antibody results in ^m6^A-Tracer enrichment around Ki-67 domains (**Fig S2B**) and yields a genome-wide domain pattern that is similar to that of the initially used antibody (**Fig S2C**). With the N-terminus antibody some of this domain pattern can also be observed, but the data quality is rather poor (**Fig S2D, E**), possibly because the antibody epitope is too far from the DNA. These results thus show that Ki-67 profiles can be reproduced with different antibodies, and are in accordance with C-terminal location of the DNA binding domain.

Finally, we sought to confirm these results by chromatin immunoprecipitation followed by sequencing (ChIP-seq). We focused on the Ki-67-AID cells, where depletion of Ki-67 by auxin treatment should cause a substantial loss of specific ChIP-seq signal. Reassuringly, when normalized over this control, cells not treated with auxin showed a domain-like ChIP-seq pattern that was very similar to the pA-DamID pattern (**Fig S3A-F**). However, the dynamic range of this ChIP-seq pattern was low compared to that of pA-DamID, and the ChIP-seq pattern was obscured when a conventional normalization over input-DNA was applied (**Fig S3C, E**), presumably because this normalization incompletely corrects for technical biases. Thus, the pA-DamID map can be recapitulated by ChIP-seq, but the signals obtained with the latter are of borderline quality. This may explain why no ChIP-seq maps have been published for Ki-67 so far.

Combined, we conclude that application of pA-DamID results in robust genome-wide Ki-67 interaction maps, although the data resolution remains limited to about 50 kb. The ^m6^A-Tracer staining indicates that interactions of Ki-67 with the genome are enriched near nucleoli, but may also occur elsewhere.

### Ki-67 binding varies between cell types, but is consistently enriched near centromeres and at small chromosomes

Nucleoli are formed around rDNA repeats that are positioned on the p-arm of several human chromosomes, next to the centromere. We therefore expected to find binding of the nucleolar protein Ki-67 near these regions. Indeed, Ki-67 interactions are enriched near centromeres of rDNA-containing chromosomes for all cell types (**Fig 1D-E**). However, Ki-67 may lack affinity for rDNA sequences themselves, as these show no enrichment in our data (**Fig S4A**). Rather, Ki-67 may interact with peri-centromeric heterochromatin (see below). Together with the ^m6^A-Tracer visualization (**Fig 1B**), these results support a model where Ki-67 interacts with chromatin at the nucleolar surface rather than in the nucleolar interior where rDNA is located (Nemeth & Grummt, 2018).

Our data indicate that Ki-67 also interacts with peri-centromeric regions of all other chromosomes (**Fig 1E-F**). We speculated that these interactions would correspond to the Ki-67 foci outside of the apparent nucleoli (**Fig 1B**). Indeed, co-immunostaining showed that centromeres (marked by CENPA) often overlap with Ki-67, even when these are not at nucleoli (**Fig 1G-H**). Moreover, the Ki-67 signal extends beyond CENPA foci (**Fig 1I**), which is in agreement with the observed binding at peri-centromeric DNA.

Differences in Ki-67 interactions among cell types are most apparent on the arms of large chromosomes (**Fig 1D, S4B**). In contrast, on small chromosomes the interactions are more consistent (**Fig S4B**), and more frequent as illustrated by a higher dynamic range compared to larger chromosomes (**Fig S4C**). Small chromosomes are typically positioned in the nuclear interior and in the vicinity of nucleoli (Bolzer *et al*, 2005; Su *et al*, 2020), suggesting that these interactions mostly involve nucleolar Ki-67.

### Release of Ki-67 from nucleoli drastically changes its DNA interactions

To test the importance of nucleolar Ki-67 for the observed genomic interactions, we released Ki-67 from nucleoli by adding a low dose of actinomycin D (ActD; 50 ng/mL) for 3 hours. ActD at this concentration specifically inhibits PolI transcription and results in nucleolar breakdown (Perry & Kelley, 1970; Ragoczy *et al.*, 2014). As a result, Ki-67 is no longer restricted to nucleoli and shows patterns that are distinct from other nucleolar markers (i.e. MKI67IP) (**Fig 2A, S5A-B**). Ki-67 localization to mitotic chromosomes is not affected by ActD (**Fig 2A**, orange arrow).

**Fig 2.**
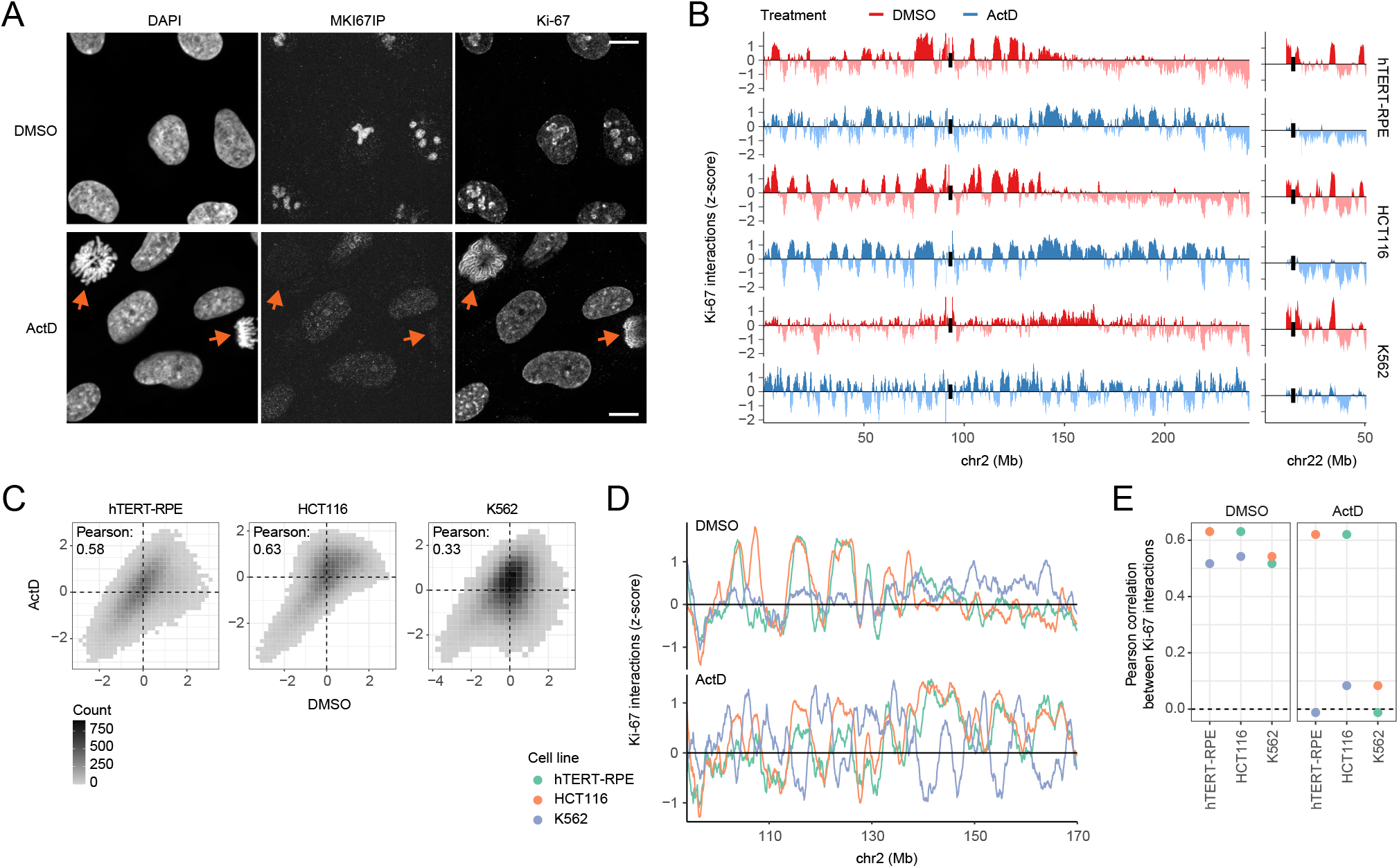
Ki-67 binding is constrained by nucleolar positioning and variable between cell types. **(A)** Representative confocal microscopy sections of hTERT-RPE cells following 3 hours of a low dose of actinomycin D (ActD, 50 ng/mL in 0.05% DMSO) to disrupt nucleoli, and of cells treated with a similar quantity of DMSO as control. Cells were stained for MKI-67IP and Ki-67 proteins to visualize nucleolar disruption and new Ki-67 localization patterns. Scale bar: 10 μm. **(B)** Comparison of Ki-67 interactions in the three cell lines after ActD-induced nucleolar disruption for two representative chromosomes. Log2-ratios were converted to z-scores to correct for differences in dynamic range between conditions and replicates. Data are averages of two experiments and smoothed with a running mean across nine 50 kb bins for visualization. Centromeres are highlighted by black bars. **(C)** Binned scatterplots between Ki-67 interactions in DMSO and ActD conditions. Pearson correlation scores were calculated with the ‘cor’ function in R. **(D)** Overlay between Ki-67 interactions between the cell types in DMSO and ActD conditions for a representative locus, as described in panel B. **(E)** Overview of Pearson correlations between Ki-67 interactions in different cell types, in DMSO and ActD conditions.

We then generated pA-DamID maps of Ki-67 (**Fig 2B**). For these quantitative analyses of Ki-67 interactions, we first converted the log_2_-ratios to z-scores. This equalizes any variable dynamic ranges between conditions and replicates without affecting the data distribution (van Schaik *et al.*, 2020). The results show that nucleolar breakdown affects Ki-67 binding to the genome in different ways depending on the cell type (**Fig 2B**). In hTERT-RPE and HCT116 cells, the genomic pattern of Ki-67 seems mostly maintained but interactions show strong quantitative differences (**Fig 2C**). This includes an overall balance shift of Ki-67 from small chromosomes to large chromosomes (**Fig S5C**). In contrast, the interaction pattern in K562 cells is more broadly altered (**Fig 2C**), although a reduction of Ki-67 interactions with rDNA-containing chromosomes is shared among all three cell types (**Fig S5C**). All three cell types also exhibit a consistent loss of Ki-67 interactions near centromeres, again most clearly for rDNA-containing chromosomes (**Fig S5D**). We verified this result by immunostaining, which showed a reduced overlap of Ki-67 with centromeres upon addition of ActD (**Fig S5E-F**). These results illustrate that nucleolar integrity is required for a normal Ki-67 interaction pattern across the genome, in particular for the preference at centromeres and small and rDNA-containing chromosomes.

### Cell cycle dynamics of Ki-67 interactions

So far, we performed these pA-DamID experiments in unsynchronized cells. However, microscopy studies have shown that Ki-67 coats chromosomes during mitosis, localizes to pre-nucleolar bodies (PNBs) in early G1 and slowly transfers to nucleoli during interphase (reviewed in Nemeth & Grummt, 2018; Verheijen *et al.*, 1989). We hypothesized that these transitions would be accompanied by a shift in genomic interactions. To test this, we synchronized RPE cells in metaphase and harvested cells at several time points during the cell cycle (van Schaik *et al.*, 2020) to profile Ki-67 interactions.

We first prepared cells for ^m6^A-Tracer microscopy to verify the localization of Ki-67 and visualize its DNA interactions. Cells synchronized in metaphase show condensed chromosomes, with Ki-67 and ^m6^A-Tracer signals at the periphery of the chromosomes (**Fig 3A**), although individual chromosomes are difficult to distinguish after pA-DamID, presumably due to several hours of nuclear permeabilization without fixation (c.f. **Fig 2A** and **3A**). One hour after synchronization, when cells have entered interphase, Ki-67 is spread throughout the nucleus in numerous PNBs (**Fig 3A**). Later time points show a gradual repositioning of Ki-67 from small PNBs to several mature nucleoli. At all interphase time points ^m6^A-Tracer signal is enriched near Ki-67 (**Fig 3A, B**), which indicates that a redistribution of Ki-67 also affects its DNA interactions.

**Fig 3.**
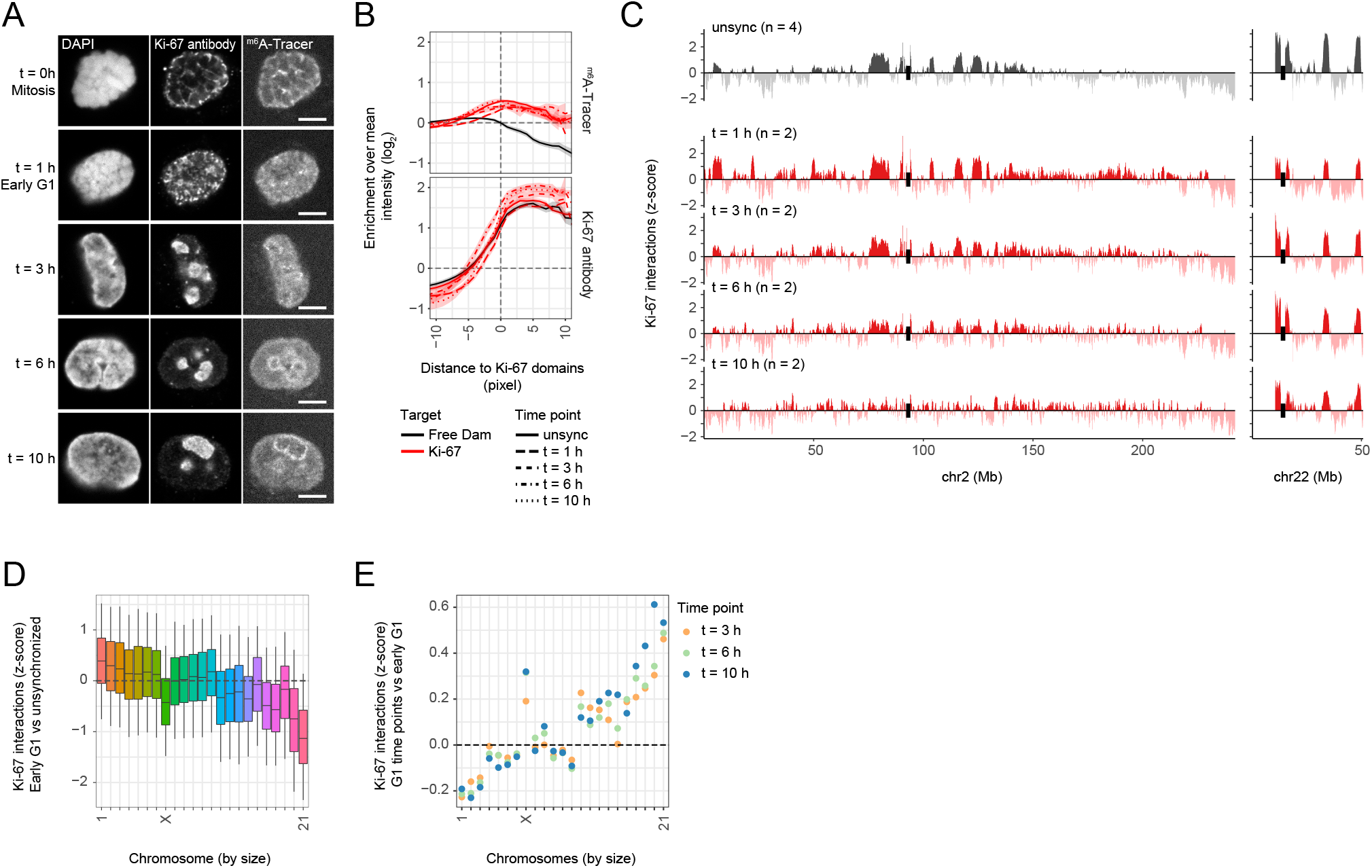
Cell cycle dynamics of Ki-67 interactions. **(A)** Representative confocal microscopy sections of hTERT-RPE cells processed with Ki-67 pA-DamID, following synchronization with a mitotic shake-off (t = 0 h, see Methods) and time points after replating of the synchronized cells. Cells were stained for Ki-67 and labeled with ^m6^A-Tracer protein. Scale bar: 5 μm. **(B)** Quantification of ^m6^A-Tracer and Ki-67 antibody enrichment near Ki-67 domains (interpreted as PNBs / nucleoli) in synchronized hTERT-RPE cells, as described in (**Fig 1C**). Instead of individual cell traces, the 95%-confidence interval of the mean is added as shaded area. Results are combined from three (unsynchronized cells) or one (synchronized cells) biological replicates. At least 10 cells were analyzed for each time point. **(C)** Ki-67 interactions profiles for the synchronized hTERT-RPE cells. Two representative chromosomes were visualized to highlight the differences between large (chr2) and small chromosomes (chr22). Data profiles are averages of *n* experiments and smoothed with a running mean across nine 50 kb bins. Centromeres are highlighted by black bars. **(D)** Chromosomal distributions of genome-wide differences between Ki-67 interactions in early G1 (t = 1 h) and unsynchronized cells. Boxplots: horizontal lines represent 25th, 50th, and 75th percentiles; whiskers extend to 5th and 95th percentiles. To test for statistical significance, a rank-based Wilcoxon test was used between chromosome size and the Ki-67 difference (p-value <2.2e-16). **(E)** Mean chromosomal differences in Ki-67 interaction scores between interphase time points and early G1 cells (t = 1 h) are plotted for chromosomes sorted by size. To test for statistical significance, Wilcoxon tests were used between chromosome size and the mean Ki-67 difference for all three comparisons (all p-values <2.2e-16).

We then processed these samples for genome-wide mapping of the ^m6^A-tagged Ki-67 interaction sites (**Fig 3C**). During mitosis (t = 0 h), Ki-67 interactions show a surprising enrichment on the distal ends of the chromosome ends (**Fig S6A, B**). However, chromosome spreads of mitotic cells processed with Ki-67 pA-DamID lack ^m6^A-Tracer signal in the middle of the mitotic cluster of chromosomes (**Fig S6C**). We therefore suspect that accessibility to the antibody and pA-Dam may be hampered by the aggregated and compacted chromosomes in mitotic cells. This accessibility bias may differentially affect distal and proximal regions of chromosomes. Indeed, ChIP-seq does not confirm the distal enrichment on mitotic chromosomes (**Fig S6D**). For these reasons, the distal enrichment as detected by pA-DamID should be interpreted with caution. Importantly, the genome-wide domain pattern that Ki-67 shows in unsynchronized cells is not visible on mitotic chromosomes, neither by pA-DamD nor by ChIP (**Fig S6A, D**), indicating that Ki-67 interacts with these domains only during interphase.

pA-DamID mapping of Ki-67 at various time points during interphase reveals striking dynamics. Early in G1 phase (t = 1 h), Ki-67 interactions are more evenly distributed among all chromosomes compared to unsynchronized cells, with increased and decreased interactions for large and small chromosomes, respectively (**Fig 3D**). Later in interphase, Ki-67 interactions are lost on large chromosomes in favor of smaller chromosomes (**Fig 3C, E**). Remarkably, chromosome X does not follow these trends. This may be caused by active repositioning of the inactive chromosome X to nucleoli during S-phase (Sun *et al*, 2017; Zhang *et al*, 2007).

These changes in genome-wide binding patterns roughly coincide with the sequential repositioning of Ki-67 from mitotic chromosomes to PNBs, and then to mature nucleoli. To further investigate the link between Ki-67 dynamics during interphase and the nucleolar maturation from PNBs, we disrupted nucleoli in hTERT-RPE cells with a short osmotic shock. Upon recovery, the process of PNB formation and nucleolar maturation is repeated (Zatsepina *et al*, 1997). Indeed, we find that Ki-67-containing PNBs are strongly enriched after 30 minutes of recovery and mostly lost again after 180 minutes (**Fig S6E**). pA-DamID maps from these cells show that the transition from the 30-minute to 180-minute time point resembles the difference between early G1 (t = 3 h) and unsynchronized cells (**Fig S6F-G**). Similar to interphase, this transition corresponds to an increase of Ki-67 interactions on small chromosomes (**Fig S6H**). We conclude that cell cycle dynamics of Ki-67 interactions correspond to a large degree to PNB maturation during interphase.

### Ki-67 and Lamin B1 interactions partially overlap and together mark the late-replicating genome

Heterochromatin positioning is redundant between nucleoli and the NL (reviewed in Bizhanova & Kaufman, 2021; Politz *et al*, 2016). To determine whether we could observe this also in our data, we compared our pA-DamID maps of Ki-67 and Lamin B1 (van Schaik *et al.*, 2020). Overall, both proteins interact with mostly similar genomic domains, although their relative interaction frequencies vary strongly (**Fig 4A, B**). Often, among the domains bound by both proteins, Ki-67 and Lamin B1 seem anti-correlated, where domains with the highest Ki-67 signals are generally lower in Lamin B1 signals, and vice-versa (**Fig 4A**, colored arrows). This partial anti-correlation is visible as the “head” of a hammer-shaped scatter plot, particularly in hTERT-RPE cells and least pronounced in K562 cells (**Fig 4C**, top panels).

**Fig 4.**
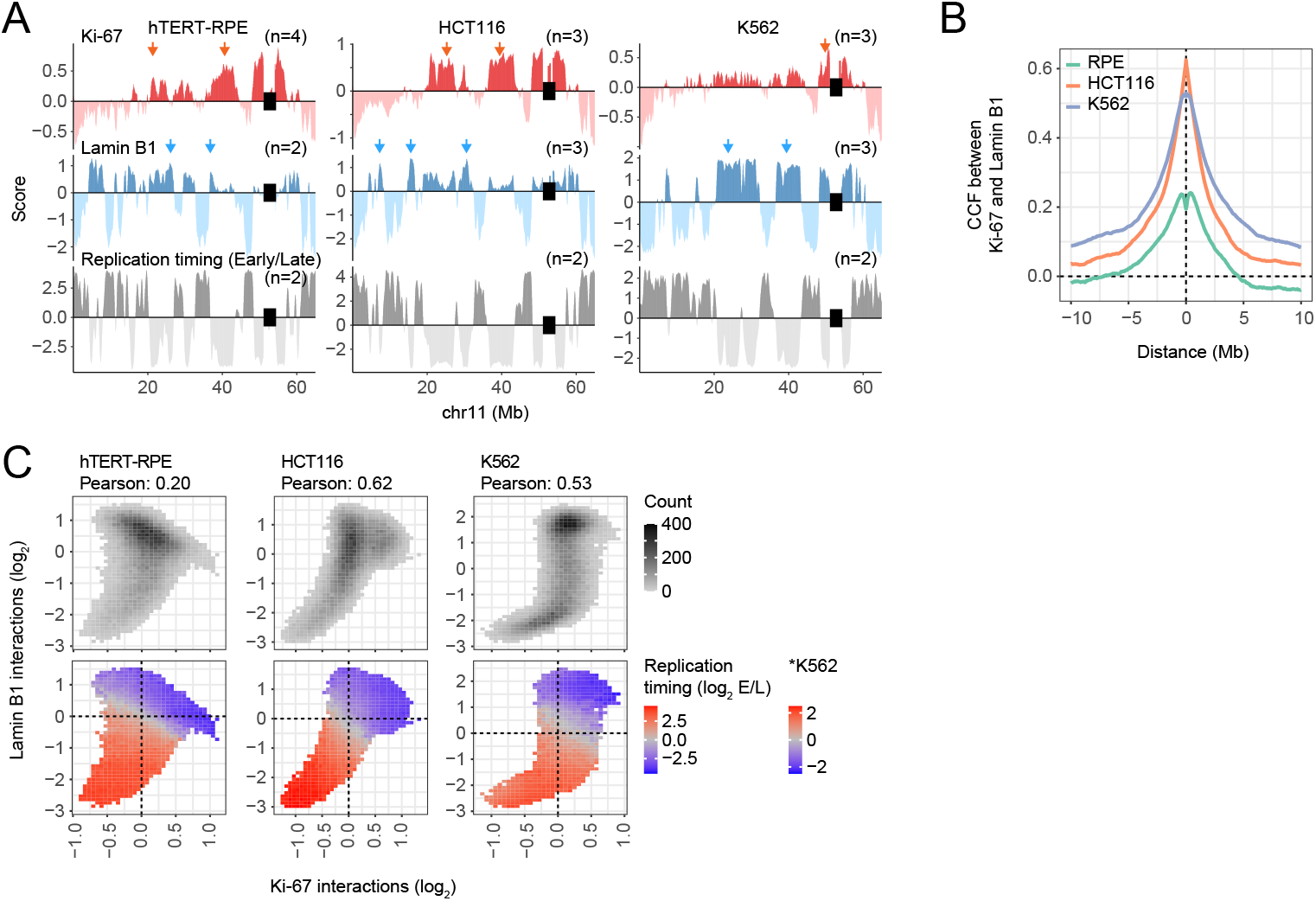
Ki-67 and Lamin B1 together mark the late-replicating genome. **(A)** Comparison of Ki-67 interactions with Lamin B1 interactions (both log_2_-ratios) and DNA replication timing (Repli-seq log_2_-ratio of Early/Late) for a representative genomic region. Arrows highlight domains with high Ki-67 and generally low Lamin B1 interactions, and vice-versa. Data profiles are averages of *n* experiments and smoothed with a running mean across nine 50 kb bins. Centromeres are highlighted by black bars. **(B)** Cross-correlation functions (CCF) between Ki-67 and Lamin B1 interactions for the three cell lines. The ‘cor.test’ function in R was used to determine statistical significance at a lag of 0, resulting in p-values <2.2e-16 for all cell lines. **(C)** Binned scatterplots between Ki-67 and Lamin B1 interactions (log_2_-ratios), showing the amount of genomic 50kb bins overlapping these points (top panels) and the mean replication timing score of overlapping genomic bins (log_2_ Early/Late) (bottom panels). Points are only shown with at least 10 overlapping genomic bins for a robust estimation of mean scores. A separate color bar is used for replication timing in K562 cells, which have a reduced dynamic range. Replication timing data were obtained from the 4D Nucleome data repository (Dekker *et al.*, 2017).

Late-replicating DNA is known to be enriched at both the nuclear periphery and nucleolus (reviewed in Marchal *et al*, 2019). Indeed, analysis of published maps of replication timing (Dekker *et al*, 2017) showed strong overlap of late-replicating domains with both the Ki-67 and Lamin B1 domains (**Fig 4A**). Strikingly, the late-replicating regions overlap almost perfectly with the hammer-heads in the Ki-67 versus Lamin B1 scatter plots (**Fig 4C**, middle panels). Thus, virtually all late-replicating domains are covered by either of the two proteins, and among these domains, competition may lead to a variable balance between the two proteins.

### Ki-67 competes with the NL for heterochromatin

To test this competition, we used the HCT116 Ki-67-AID cell line and mapped Lamin B1 interactions following 24 hours of Ki-67 depletion. Additionally, given the roles of Ki-67 in mitosis and the reorganization of the genome afterwards, we combined Ki-67 depletion with two cell synchronization strategies. First, Ki-67 was depleted during a double thymidine block. These cells undergo a single cell division and the synchronization prevents any cell cycle perturbation that Ki-67 depletion may induce. Second, to specifically test whether mitosis is required for DNA repositioning, Ki-67 was depleted in G2-arrested cells using a CDK1 inhibitor (**Fig 5A**). Flow cytometry confirmed cell synchronization with and without Ki-67 depletion (**Fig 5A**, bottom panels, **S7A**). We first verified the Ki-67 pA-DamID patterns in synchronized cells with normal Ki-67 levels (i.e., not treated with auxin). Here, we found a shift of Ki-67 between S-phase and G2 cells that correlates with chromosome size (**Fig 5B, S7B**), akin to the progressive shift towards smaller chromosomes that we observed during interphase in hTERT-RPE cells (**Fig 3C, E**).

**Fig 5.**
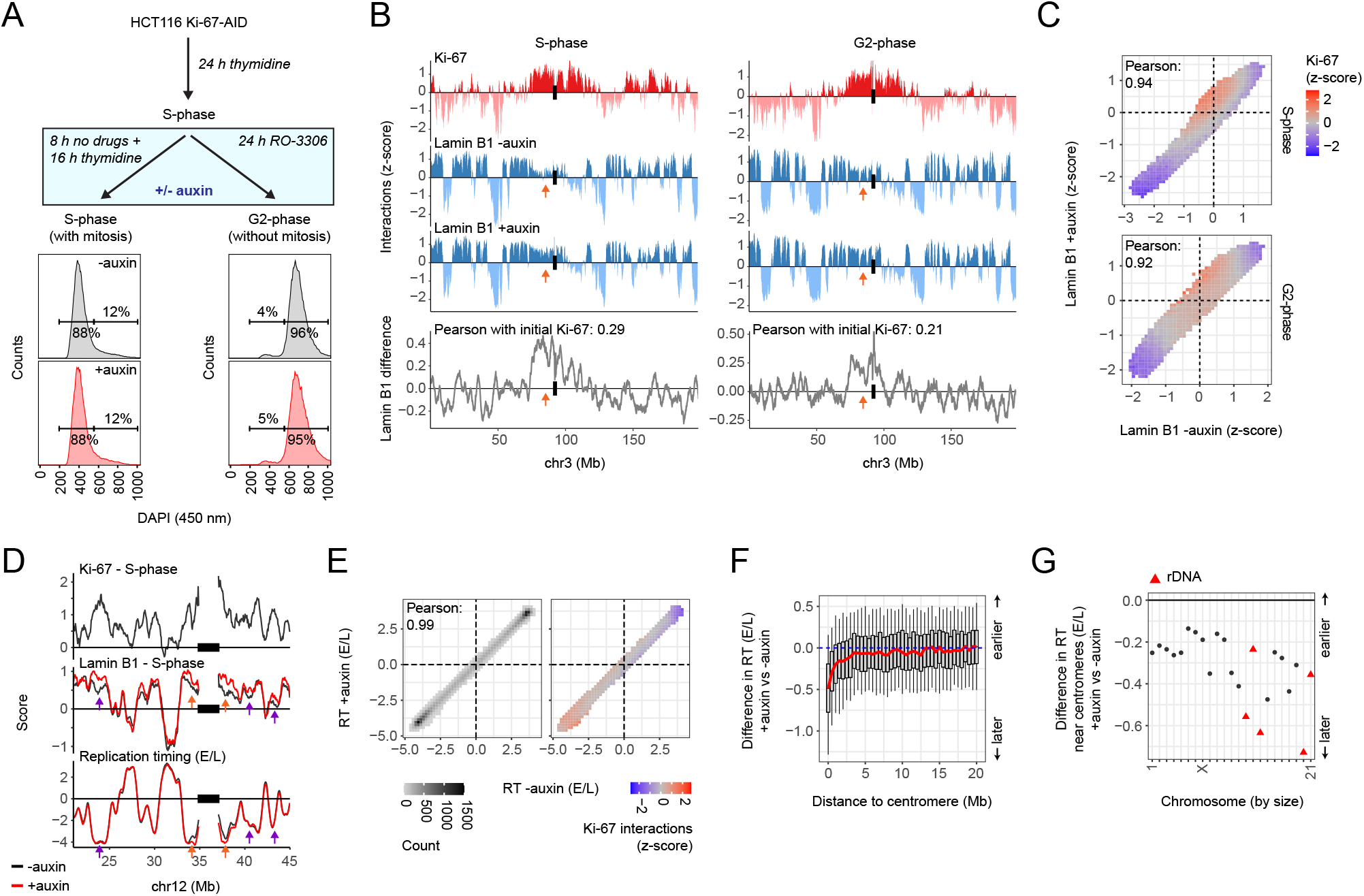
Effects of Ki-67 depletion on Lamin B1 interactions and replication timing. **(A)** Overview of the Ki-67 depletion experiment in HCT116 Ki-67-AID cells (top). Cells were either cultured with a double thymidine arrest to enrich for S-phase cells (which includes one cell division) or arrested in G2 by CDK1 inhibition immediately following an initial S-phase arrest (which prevents a cell division). FACS analysis into G1/S and G2 populations confirmed successful synchronization and illustrates that Ki-67 depletion does not affect this (bottom panels). **(B)** Representative genomic region showing the effect of Ki-67 depletion on Lamin B1 interactions, for HCT116 Ki-67-AID cells synchronized in S-phase (left panel) and G2-phase (right panel). Ki-67 interactions were only profiled at non-depleted levels. Orange arrows highlight regions with high Ki-67 interactions that gain Lamin B1 interactions upon Ki-67 depletion. Data profiles are averages of 2 experiments and smoothed with a running mean. The centromere is located just to the right of the depicted region. **(C)** Binned scatterplots of Lamin B1 interactions at in HCT116 cells with normal versus depleted Ki-67 levels, as described in (**Fig 4C**). Bins are colored by mean Ki-67 interactions in S-phase (top panels) and G2-phase (bottom panels). Signals are smoothed with a running mean of three 50 kb bins to reduce noise in the data. **(D)** Representative genomic region showing the effect of auxin-mediated Ki-67 depletion on (S-phase) Lamin B1 interactions and replication timing (log2 Early/Late) in HCT116 Ki-67-AID cells, as described in panel B. Orange arrows highlight domains with gained Lamin B1 interactions and delayed replication timing; purple arrows highlight domains with gained Lamin B1 interactions but unaffected replication timing. **(E)** Binned scatterplots (as in panel C) of replication timing (RT, E/L) in cells with normal versus depleted Ki-67 levels, colored by the density of overlapping 50 kb bins (left panel) and mean Ki-67 interactions at normal Ki-67 levels (right panel). **(F)** Distribution of differences in replication timing (RT) nearby centromeres. Boxplots: horizontal lines represent 25th, 50th, and 75th percentiles; whiskers extend to 5th and 95th percentiles. **(G)** Mean difference in replication timing (RT) within 2 Mb from the centromere of each chromosome. rDNA-containing chromosomes are highlighted in red.

Ki-67 depletion induces small quantitative differences in Lamin B1 interactions (**Fig 5B**, orange arrows). Importantly, increases in Lamin B1 interactions are most prominent in regions that were originally bound by Ki-67 (**Fig 5C**). This effect is most apparent in S-phase cells (which had progressed through mitosis), but is still present in cells blocked in G2 (with Pearson correlations between the initial Ki-67 score and Lamin B1 difference of approximately 0.3 and 0.2 for the two conditions, respectively). Passage through mitosis is thus not required to change genome positioning between Ki-67 and the NL. We conclude that Ki-67 counteracts Lamin B1 interactions of a subset of late-replicating domains.

### Ki-67 does not detectably control 3D genome interactions in interphase

Considering the effects of Ki-67 on NL interactions, we wondered whether it also controls other aspects of 3D genome organization. To test this, we performed Hi-C experiments in HCT116 Ki-67-AID cells that were synchronized in S-phase (**Fig 5A**). Remarkably, depletion of Ki-67 does not affect genome interactions in any aspect that we explored, including compartment A/B formation and overall folding of the chromosomes (**Fig S8A-I**). In particular, the compartment scores across the genome are virtually identical between the + and - auxin conditions, and any subtle change of these scores is not linked to the level of Ki-67 binding (**Fig S8F**) nor to linear distance from centromeres (**Fig S8G**). Combined, these experiments indicate that Ki-67 modulates positioning of genomic regions relative to the NL, without any detectable effect on chromatin–chromatin interactions.

For comparison, we also generated Hi-C data in cells synchronized in metaphase. Here, we did observe that depletion of Ki-67 increases inter-chromosomal interactions and very long-range cis-interactions (>10 Mb) (**Fig S8A-C**). In particular, Ki-67 seems important to prevent the chromosome arms from folding back onto each other (**Fig S8J**). This result is in accordance with the model that Ki-67 acts as a surfactant that prevents intermingling of mitotic chromosomes (Cuylen *et al.*, 2016), and underscores that the lack of detectable effects of Ki-67 depletion on 3D genome interactions is not due to technical issues with the Hi-C data.

### Ki-67 promotes timely replication of centromeres

Next, we asked whether Ki-67 depletion affected the replication timing of the regions that it interacts with. We tested this by Repli-seq (Marchal *et al*, 2018), 24 hours after Ki-67 depletion in (unsynchronized) HCT116 cells (**Fig 5D, E**). While replication timing across most of the genome remained unaffected by this depletion, we did observe a small but consistent delay in replication timing in peri-centromeric regions (**Fig 5F, G**). Centromeric sequences, which cannot be uniquely mapped to individual chromosomes, also showed a delayed replication timing upon Ki-67 depletion (**Fig S7C**). The replication timing observations are different from Lamin B1 interactions that increased near centromeres but also on other domains with high Ki-67 scores (**Fig 5D,** purple arrows, **S7D, E**). Thus, besides its known roles in mitosis, Ki-67 specifically affects the replication timing of (peri-)centromeric regions in interphase.

### No evidence for direct gene repression by Ki-67

Genes in late replicating and NL-associated domains generally show little transcriptional activity (Guelen *et al.*, 2008; Hiratani *et al*, 2008; reviewed in Marchal *et al.*, 2019; Reddy *et al*, 2008; van Steensel & Belmont, 2017). Indeed, mRNA-seq data show that genes in the above-mentioned scatterplot hammer-head predominantly are lowly transcribed (**Fig S7F**). In line with this, genes with high levels of Ki-67 tend to be inactive, particularly in hTERT-RPE and HTC116 cells.

We then asked whether Ki-67 contributes directly to the transcriptional repression of these genes. Several studies have found changes in gene activity upon Ki-67 loss (Garwain *et al*, 2020; Mrouj *et al*, 2021; Sobecki *et al.*, 2016; Sun *et al.*, 2017), but none could attribute this to direct effects, in part because the binding pattern of Ki-67 was not known. We therefore generated RNA-seq data in the HCT116 Ki-67-AID cells. We again synchronized these cells in S-phase, to rule out any cell cycle effects that Ki-67 depletion may have in these cells (**Fig 5A**). Intriguingly, we find that gene expression is largely unaffected by acute Ki-67 depletion, with only three genes reaching our significance cut-off: ALDH1A3, CYP1A1 and HAS3 (**Fig S7G**). We cannot link these genes to initial Ki-67 binding or Ki-67 biology. A similar result was obtained in a recent study that also employed auxin-mediated Ki-67-AID depletion in HCT116 cells (Garwain *et al.*, 2020).

We also compared Ki-67 interactions with gene expression data following Ki-67 depletion in hTERT-RPE cells (Sun *et al.*, 2017). Unlike HTC116, these cells express p21, which causes cell cycle perturbations in response to Ki-67 loss, and concomitant differential expression of thousands of genes (Sun *et al.*, 2017). Our re-analysis of these data indicates that the changes in gene expression do not correlate with initial Ki-67 binding (**Fig S7H**). This further indicates that Ki-67 binding is not directly involved in gene silencing.

### Ki-67 binding correlates positively with H3K9me3 and negatively with H3K27me3

Finally, we compared the Ki-67 patterns to those of the main repressive histone modifications H3K27me3 and H3K9me3 (**Fig 6A**). Remarkably, these marks are strongly correlated with the balance between Ki-67 and Lamin B1. In hTERT-RPE and HCT116 cells the scatterplot hammer-head shows a strong partitioning, with Ki-67 interactions being positively linked to H3K9me3 but not H3K27me3 (**Fig 6B**). In K562 cells the preference of H3K9me3 for Ki-67-rich regions is less pronounced, although this may be due to a lower signal/noise ratio of the Ki-67 pA-DamID map in this cell line.

**Fig 6.**
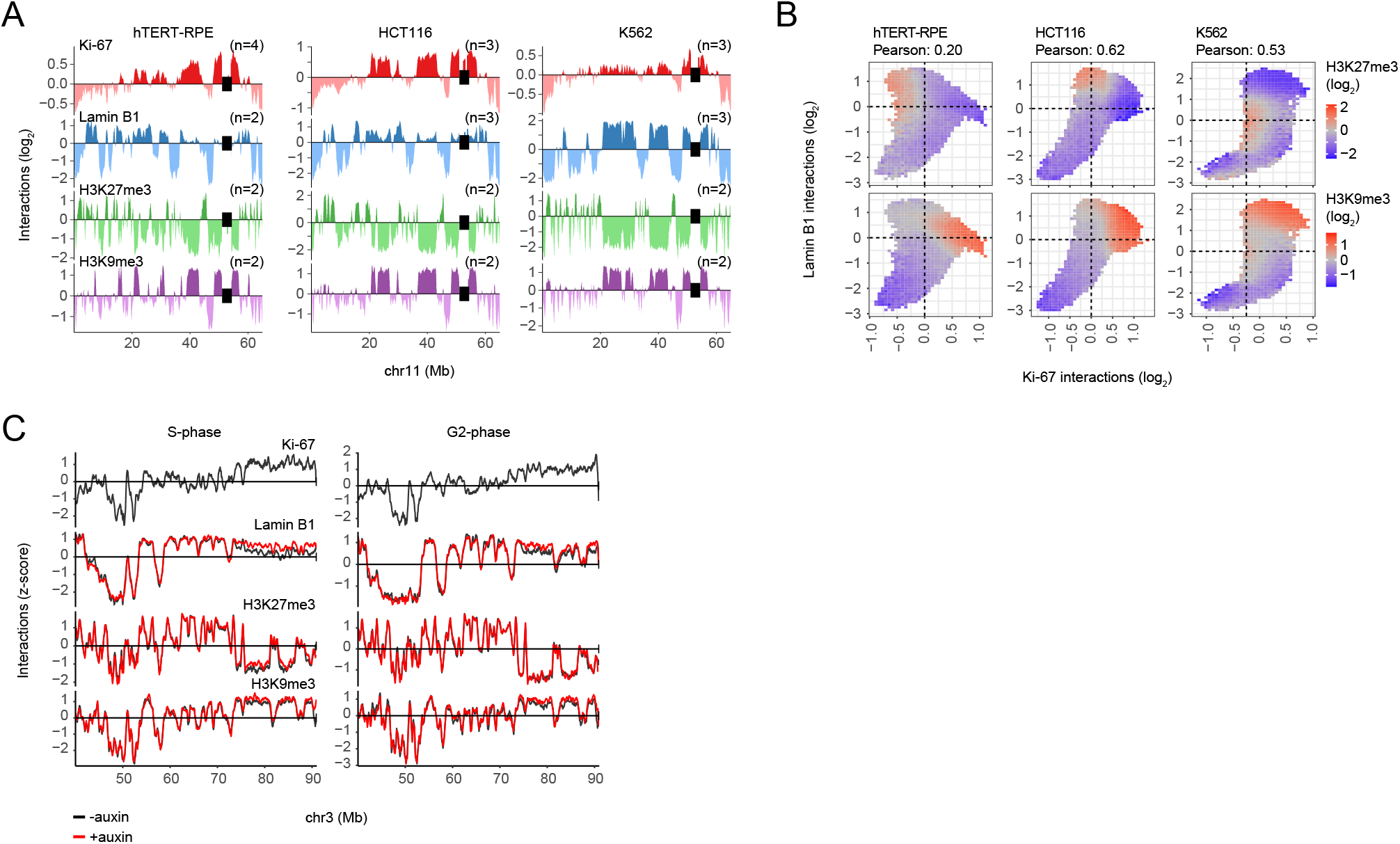
Comparison of the Ki-67 and Lamin B1 balance with H3K27me3 and H3K9me3. **(A)** Comparison of Ki-67 interactions with Lamin B1 interactions and H3K27me3 and H3K9me3 histone modifications for a representative genomic region. Data profiles are averages of *n* pA-DamID experiments and smoothed with a running mean across 9 50kb bins. Centromeres are highlighted by black bars. **(B)** Binned scatterplots of Ki-67 and Lamin B1 interactions as in (**Fig 4C**), colored according to the mean H3K27me3 and H3K9me3 signals. **(C)** A similar figure as (**Fig 5D**), but additionally showing the effect of Ki-67 depletion on H3K27me3 and H3K9me3.

Following acute Ki-67 depletion in HCT116 cells (**Fig 5**), H3K27me3 and H3K9me3 are only subtly affected, with effect sizes that seem smaller than we observed for Lamin B1 interactions (**Fig 6C**). Ki-67 binding is thus correlated with the epigenetic makeup of chromatin, but not required to maintain it.

## DISCUSSION

Microscopy studies have illustrated that Ki-67 and nucleoli are important contributors to genome organization, in particular for heterochromatin positioning and the separation of mitotic chromosomes (Booth *et al.*, 2014; Cuylen *et al.*, 2016; Cuylen-Haering *et al.*, 2020; Sobecki *et al.*, 2016; Takagi *et al.*, 2016). However, a lack of genome-wide interaction profiles of Ki-67 has hampered a detailed understanding of how this protein shapes and regulates the genome. In this work, we applied pA-DamID to profile genome - Ki-67 interactions in several human cell lines and throughout interphase. In our hands, pA-DamID appears to be more suited to study K-67 than ChIP-seq.

Early after mitosis, we observed a rapid rearrangement of Ki-67-DNA interactions across the genome. Within ~1 hour, Ki-67 moves to many late-replicating domains. This transition may be triggered by dephosphorylation of Ki-67 (MacCallum & Hall, 1999; Saiwaki *et al*, 2005) (Yamazaki *et al*, 2022). Our data are in accordance with early microscopy observations that Ki-67 overlaps with most large heterochromatin domains in early interphase (Bridger *et al*, 1998). However, our results also show that Ki-67 gradually redistributes during early G1 from large to smaller chromosomes. This redistribution roughly coincides with the relocation of Ki-67 from PNBs to nucleoli, and is reverted by perturbations that release Ki-67 from nucleoli and re-establish PNBs. We therefore propose that Ki-67 in PNBs interacts with late-replicating domains throughout the genome, while upon sequestration of Ki-67 at nucleoli these interactions become restricted to small chromosomes. In support of this model, small chromosomes tend to be positioned near nucleoli (Quinodoz *et al*, 2018; Su *et al.*, 2020).

Our results implicate Ki-67 as one of the proteins that modulate the distribution of late-replicating DNA between nucleoli and the NL (reviewed in Bizhanova & Kaufman, 2021; Politz *et al.*, 2016; van Steensel & Belmont, 2017). Most likely this is due to direct interaction of Ki-67 with a subset of late-replicating regions. It is interesting to note that both Ki-67 and lamins preferentially bind AT-rich DNA in vitro (Luderus *et al*, 1994; MacCallum & Hall, 2000), which matches observations that both nucleolus- and lamina-associated domains are AT-rich (Meuleman *et al*, 2013; van Koningsbruggen *et al.*, 2010). However, it is likely that other proteins besides Ki-67 mediate nucleolar interactions. For example, different from our Ki-67 profiles, a subset of nucleolar interactions shows enrichment for H3K27me3 and early-replicating DNA (Vertii *et al.*, 2019). Moreover, Ki-67 is slowly degraded in G1 and completely absent in senescent cells (Gerdes *et al*, 1984; Miller *et al*, 2018; Sobecki *et al*,2017), yet these cells largely maintain their nucleolus - genome interactions (Dillinger *et al.*,2017). Exceptions are alpha-satellite sequences (at centromeres) and H3K9me3-marked chromatin, which lose nucleolar positioning upon senescence (Dillinger *et al.*, 2017). Our data show that these regions strongly interact with Ki-67, supporting a direct role in nucleolus tethering.

Surprisingly, this tethering does not have any detectable impact on A/B compartmentalization as detected by Hi-C. Thus, NL interactions and A/B compartmentalization can be uncoupled, and Ki-67 controls the former but not the latter.

This result is reminiscent of Hi-C experiments in cells devoid of both Lamin A and LBR. These cells show a partially inverted organization with active chromatin at the nuclear periphery and nearly all heterochromatin in the nuclear interior, while A/B compartmentalization is largely unaffected (Falk *et al*, 2019). Redistribution of heterochromatin between nucleoli and the NL can thus occur independently of 3D genome interactions.

Upon Ki-67 depletion, we observed a surprising delay in replication timing specifically at (peri-)centromeres. These regions show increased Lamin B1 interactions, yet other regions that reposition towards the NL are not affected in their replication. Possibly, replication timing of centromeric regions is more sensitive to the positioning relative to the NL than that of other genomic loci. Alternatively Ki-67 may directly affect replication of centromeres, which are known for their middle-late and tightly regulated DNA replication (reviewed in Barra & Fachinetti, 2018; Bloom & Costanzo, 2017). The perturbed replication may be consequence of increased transcription of alpha-satellite sequences following Ki-67 depletion (Bury *et al*, 2020). Resulting transcription-replication conflicts could result in delayed and incomplete DNA replication, as was recently shown following rapid CENP-A depletion (Giunta *et al*, 2021).

One of the fascinating aspects of Ki-67 is its role on mitotic chromosomes, where it establishes the peri-chromosomal layer (Booth *et al.*, 2014; Gerdes *et al.*, 1984; Stenstrom *et al*, 2020; Verheijen *et al.*, 1989) and acts as a surfactant that prevents chromosomal intermingling (Cuylen *et al.*, 2016; Takagi *et al.*, 2016). In accordance with this role, our mitotic Hi-C data suggest that Ki-67 suppresses trans-interactions and very long-range cis-interactions. We emphasize that this result needs to be further confirmed by additional experiments. Furthermore, pA-DamID and ChIP-seq indicate that mitotic Ki-67 does not associate with large domains as is observed in interphase. It is known that mitotic Ki-67 has reduced DNA binding affinity (MacCallum & Hall, 1999), although the mechanism remains unclear.

Ki-67 is an important protein for faithful chromosome segregation. It is degraded following mitosis in G1, but protein levels accumulate again in early S-phase (Miller *et al.*,2018; Sobecki *et al.*, 2017). It has been unclear whether this early accumulation was required to build up sufficient protein levels for mitosis, or whether Ki-67 also has a functional role in interphase. Here, we provided support for the latter hypothesis, given that Ki-67 controls the nucleolus - NL balance of interphase heterochromatin and is required for timely replication of centromeres.

## METHODS

### Experimental methods

#### Cell culture

hTERT-RPE cells (ATCC CRL-4000), HCT116 cells (ATCC CCL-247) and K562 cells (ATCC CCL-243) were cultured according to their ATCC handling procedure, except that cells were passaged using 0.05%Trypsin-EDTA (Gibco) instead of 0.25%. Adherent cells (hTERT-RPE cells and HCT116 cells) were harvested with 0.05% Trypsin-EDTA, which was inactivated with complete medium.

HCT116 Ki-67-miniAID-mClover cells (Ki-67-AID in short) were kindly provided by M. Takagi and cultured as wildtype HCT116 cells. Ki-67 depletion was induced by adding 500 μM auxin for 24 hours (from a 1 mM stock solution) (Sigma-Aldrich), or a similar amount of DMSO for control cells.

All cells were tested for mycoplasma presence every 2 months.

#### Cell cycle synchronization

Cell cycle synchronization of hTERT-RPE was performed as described previously (van Schaik *et al.*, 2020). Cells were cultured for 24 h in 75 cm^2^ flasks to reach ~50% confluency, after which mitotic cells were enriched for using a sequential synchronization strategy of G2 arrest (16 h of 10 μM RO-3306, CDK1 inhibitor) followed by metaphase blocking (1.5 h of 25 ng/mL nocodazole, microtubule inhibitor). Mitotic cells were harvested by shake-off and immediately processed for pA-DamID or replated in complete medium for interphase time points. Light microscopy confirmed successful mitotic synchronization, and adherence and cell division of the replated cells. Replated cells were harvested after 1, 3, 6 and 10 h. A similar approach was used to synchronize HCT116 Ki-67-AID cells in metaphase, except that 8 μM of RO-3306 was used to arrest the cells in G2.

#### Osmotic shock

hTERT-RPE cells were cultured overnight in multiple 10 cm dishes with 10 mm cover slips. Cells were washed in pre-warmed 1x phosphate-buffer saline (PBS), after which nucleoli were disrupted by a 15-minute osmotic shock (0.2x PBS) (Zatsepina *et al.*, 1997). Cells were released in complete medium and harvested for pA-DamID after 30 and 180 minutes. To verify nucleolar reassembly, cover slips with attached cells were removed before harvesting, washed in PBS and fixed for 10 minutes in 2% formaldehyde/PBS.

#### Actinomycin D treatment

hTERT-RPE cells, HCT116 cells and K562 cells were cultured overnight in two 10 cm dishes with 10 mm cover slips. PolI transcription was specifically inhibited using a 3 h treatment of 50 ng/mL actinomycin D (from a 1 mg/mL stock solution). A similar amount of DMSO was added to control cells. Cover slips were removed and cells were fixed with 2% formaldehyde/PBS to verify nucleolar disruption and Ki-67 redistribution. For K562 cells, the cover slips were coated with poly-L-lysine to immobilize the cells. Remaining cells were harvested for pA-DamID experiments.

#### HCT116 Ki-67-AID synchronization experiments

HCT116 Ki-67-AID (M. Takagi) were synchronized using a sequential arrest into S-phase and G2-phase. First, cells were cultured overnight in 2.5 mM thymidine to block cells in S-phase, and then washed two times in pre-warmed PBS and once in complete medium. For S-phase cells, the cells were released in complete medium for 8 h, after which they were incubated with 2.5 mM thymidine for another 16 h. For G2-phase cells, the cells were released on complete medium supplemented with 10 μM RO-3306 (CDK1 inhibitor) for 24 h. During the last 24 h, Ki-67 depletion was induced by adding 500 μM auxin (Sigma-Aldrich), or a similar amount of DMSO for control cells. For all conditions, cells were cultured in 10 cm dishes with 10 mm poly-L-lysine coated coverslips. Before harvesting cells for pA-DamID, cover slips were removed, washed in PBS and cells were fixed for 10 minutes with 2% formaldehyde/PBS to verify Ki-67 depletion. Additionally, cells synchronization was confirmed using ethanol fixation and DNA staining with 1 μg/mL DAPI (Marchal *et al.*, 2018). DNA content was measured using flow cytometry on an Attune NxT (Invitrogen, Thermo Fisher Scientific).

#### pA-DamID

pA-DamID experiments were performed as described previously (van Schaik *et al.*, 2020). One million cells were collected by centrifugation in 1.5 mL Eppendorf tubes (3 minutes, 500g, at 4°C), washed in 400 μL of ice-cold PBS and subsequently washed in 400 μL of ice-cold digitonin wash buffer (DigWash) (20 mM HEPES-KOH pH 7.5, 150 mM NaCl, 0.5 mM spermidine, 0.02% digitonin, cOmplete Protease Inhibitor Cocktail (Roche # 11873580001)). Cells were resuspended in 200 μL DigWash supplemented with a primary antibody and rotated for 2 hours at 4 °C. After a DigWash wash step, cells were resuspended in 200 μL DigWash supplemented with 1:200 pA-Dam protein (~40 units of Dam activity) and rotated for 1 hour at 4 °C. After two wash steps with DigWash, Dam activity was induced by resuspension in 100 μL DigWash supplemented with 80 μM S-adenosyl-methionine (SAM) (NEB, B9003S) and incubated at 37 °C for 30 minutes. For every experimental condition, one million cells were also processed without antibodies or pA-Dam protein, but instead incubated with 4 units of Dam enzyme (NEB, M0222L) during the activating step. This freely diffusing Dam moiety serves as control to account for DNA accessibility and amplification biases.

For the primary antibody incubation, we used 1:100 dilutions for antibodies against Ki-67 (Abcam ab15580, rabbit), N-terminal Ki-67 (Atlas Antibodies HPA001164-25UL, rabbit), C-terminal Ki-67 (Novus NB600-1252-0.1ml, rabbit), H3K27me3 (CST C36B11, rabbit), and H3K9me3 (Diagenode C15410193, rabbit). A 1:400 dilution was used for the antibody against Lamin B1 (Abcam ab16048, rabbit).

At this point, either genomic DNA was isolated and processed for high-throughput sequencing, or cells were diluted in PBS and plated on poly-L-lysine coated cover slips for 30 minutes, followed by 10 minutes fixation with 2% formaldehyde/PBS and immunostaining. These cover slips were first coated with 0.1% (w/v) poly-L-lysine (Sigma-Aldrich, #P8920) for 15 minutes, washed with H_2_O (once) and PBS (3 times) and stored in 70% ethanol for later use. Samples were prepared for single-end 65 bps Illumina Hi-seq sequencing as described previously (Leemans *et al*, 2019; Vogel *et al*, 2007), except that the DpnII digestion was omitted. Approximately 15 million reads were sequenced for every Ki-67 and Dam control sample to ensure high quality Ki-67 interaction data, and approximately 5 million reads were sequenced for Lamin B1, H3K27me3 and H3K9me3 libraries.

#### Repli-seq

Following 24 h of Ki-67 depletion in HCT116 Ki-67-AID cells (see above), BrdU was added to a final concentration of 100 μM and cells were incubated at 37 °C for another 2 h. Cells were fixed in ice-cold ethanol and processed for Repli-seq as described (Marchal *et al.*,2018). Libraries were sequenced with approximately 10 million 2×150 bps paired-end reads per sample on an Illumina NovaSeq 6000.

#### RNA-seq

HCT116 Ki-67-AID cells were synchronized in S-phase with 24 h of Ki-67 depletion (see above). RNA was isolated using the Qiagen RNeasy column purification kit, after which sequencing libraries were prepared using the Illumina TruSeq polyA stranded RNA kit. Libraries were sequenced with approximately 25 million 2×54 bps paired-end reads on an Illumina NovaSeq 6000.

#### ChIP-seq

Unsynchronized and metaphase-synchronized HCT116 Ki-67-AID cells were treated for 24 h with auxin to induce Ki-67 depletion, or an equivalent amount of DMSO only (see above). Cells were fixed in 1% formaldehyde for 10 minutes after which the reaction was quenched with 2.0 M glycine and the cells were lysed. A Bioruptor Plus sonication device (Diagenode) was used to fragment chromatin to ~300 bps fragments. Ki-67 antibody (Abcam ab15580, 5 μL per ChIP) was coupled to Protein G beads (Thermo Fisher Scientific), and incubated overnight (4 °C) with the chromatin fragments. The chromatin was then washed, eluted from the beads, reverse crosslinked and purified with the MinElute PCR purification kit (Qiagen). The DNA fragments were prepared with the KAPA HTP Library Preparation Kit (Roche) for sequencing. Libraries were sequenced with approximately 30 million 75 bps single-end reads per sample on an Illumina NextSeq.

#### Hi-C

S-phase HCT116 Ki-67-AID cells treated with and without addition of auxin to deplete Ki-67 (see above) were processed for Hi-C as previously described (Haarhuis *et al*, 2017) with minor modifications (Liu *et al*, 2021). For each sample, 10 million cells were harvested and cross-linked with 2% formaldehyde. After digestion with MboI in the nucleus, biotinylated nucleotides were incorporated at the restriction overhangs, joined by blunt-end ligation, and used to enriched by streptavidin pulldown. Libraries for sequencing were prepared using a standard end-repair and 3A-tailing method. Libraries were sequenced with approximately 100 million 2×50 bps paired-end reads per sample on an Illumina NovaSeq 6000.

For Hi-C on the mitotic cells, which lack intact nuclei, a different digestion strategy was used to maintain cellular integrity (Elbatsh *et al*, 2019). Following centrifugation (5 minutes, 1000g), the pellet was resuspended in 500 μL 1x CutSmart buffer. 5 μL of 10% SDS was carefully mixed by pipetting up and down, and incubated for 10 minutes at 65 °C. Next, 30 μL of 20% Triton was added in the same way, before addition of 40 μL of MboI (5 U / μL) and overnight incubation at 37 °C.

#### Immunostaining and microscopy

Cover slips with formaldehyde fixed cells were washed with PBS, permeabilized with 0.5% NP40/PBS for 20 minutes and blocked with 1% BSA/PBS for 1 h. Cover slips were incubated with primary antibodies at room temperature for 1 h, and washed three times with PBS. Next, cover slips were incubated with secondary antibodies at room temperature for 45 minutes. After one wash with PBS, DNA was stained with 1 μg/mL DAPI for 10 minutes, followed by two washes with PBS and one with H_2_O. Cover slips were mounted with Vectashield and sealed with nail polish.

Cells processed with Ki-67 pA-DamID were first incubated with an antibody against Ki-67 (1:1000, Abcam ab15580, rabbit), and then with ^m6^A-Tracer protein (1:500, 1.15 mg/mL) and an Alexa594-fused antibody against rabbit (1:250, Jackson 711-585-152, donkey). Cells that received the osmotic shock or actinomycin D were first incubated with antibodies against MKI-67IP (1:500, Atlas Antibodies AMAB90961, mouse) and Ki-67 (1:1000, Abcam ab15580, rabbit), and then with an Alexa488-fused antibody against mouse (1:1000, ThermoFisher A11001, goat) and an Alexa594-fused antibody against rabbit (1:250, Jackson 711-585-152, donkey). HCT116 Ki-67-miniAID-mClover cells were first incubated with an antibody against Ki-67 (1:1000, Abcam ab15580, rabbit), and then an Alexa594-fused antibody against rabbit (1:250, Jackson 711-585-152, donkey). Wildtype hTERT-RPE cells were stained with antibodies against CENPA (1:500, Abcam ab13939, mouse) and Ki-67 (1:1000, Abcam ab15580, rabbit), and then with an Alexa488-fused antibody against mouse (1:1000, ThermoFisher A11001, goat) and an Alexa594-fused antibody against rabbit (1:250, Jackson 711-585-152, donkey).

To prepare chromosome spreads of metaphase HCT116 Ki-67-AID cells processed with pA-DamID using a Ki-67 antibody or as Dam control, the cells were first incubated in hypotonic conditions (75 mM KCl at 37 °C for 10 minutes). Afterwards, chromosomes were spread on a 12 mm cover slip using centrifugation by cytospin (800 RPM for 3 minutes), immediately followed by fixation (2% formaldehyde for 10 minutes). Immunostaining was performed as described above.

Single 1024×1024-pixel confocal sections around the middle plane of nuclei were imaged on a Leica SP5 with a 63× NA 1.47 oil immersion objective, using bidirectional scanning, and 8× line averaging. An additional 3× electronic zoom was used for images that required a higher magnification. Alternatively, z-stacks were imaged using 2x line averaging and a 380 nm step size.

### Computational analyses

#### pA-DamID processing

pA-DamID reads were filtered to contain the DpnI adapter sequence, which was then trimmed off with cutadapt 1.11 and custom scripts. The remaining genomic DNA sequence was mapped to a hybrid genome consisting of GRCh38 v15 (without alternative haplotypes) and a ribosomal model (GenBank: U13369.1) with bwa mem 0.7.17. Further data processing was performed with custom R scripts. Reads were filtered to have a mapping quality of at least 10 and overlap with the ends of a GATC fragment, and subsequently counted in genomic bins of different sizes. In this manuscript, 50 kb genomic bins were used as a compromise between resolution and reproducibility of Ki-67 interaction profiles.

Counts were normalized to 1 million reads and a log_2_-ratio was calculated over the Dam control sample. At least two biological replicates were generated for each condition, which ratios were averaged for downstream analyses. For quantitative comparisons, the log_2_-ratios were converted to z-scores to account for differences in dynamic range between conditions and replicates (van Schaik *et al.*, 2020).

#### Repli-seq

Adapters and low-quality reads were removed from Repli-seq reads with fastp 0.12.2 (Chen *et al*, 2018). Trimmed reads were mapped and processed as pA-DamID reads with several differences. First, duplicated reads were removed. Second, reads were not required to overlap with the ends of GATC fragments. Third, log_2_-ratios of early/late were quantile normalized instead of z-score normalization.

#### RNA-seq

Adapter sequences and low-quality reads were removed from RNA-seq reads with fastp 0.12.2 (Chen *et al.*, 2018). Trimmed reads were aligned to GRCh38 v15 and counted in gencode v24 genes with STAR 2.5.4a (Dobin *et al*, 2013). DESeq2 1.30.1 was used to call differentially expressed genes (Love *et al*, 2014), which we defined as genes that have a Benjamini-Hochberg corrected p-value lower than 0.05 when testing for a log_2_-fold change different than 0. FPKM values were calculated with ‘fpkm()’ in DESeq2 using the combined exon length as gene length. The mean FPKM value between replicates was used for downstream analyses.

#### ChIP-seq

ChIP-seq reads were processed similar to the Repli-seq reads as described above, except that log_2_-ratios of Ki-67 antibody / input were not quantile-normalized but z-transformed similar to the pA-DamID data.

#### Hi-C

Hi-C reads were processed with HiC-Pro (Servant *et al*, 2015), which performs mapping, generation of contact matrices (100 kb, 1 Mb) and ICE normalization. Matrices were loaded in R and further processed with GENOVA 1.0.0.9 (van der Weide *et al*, 2021).

#### Image analysis

Image analysis was performed with custom ImageJ 2.0.0 and R scripts.

To calculate ^m6^A-Tracer enrichment around nucleoli, individual nuclei were first segmented using the DAPI signal. This was done using a sequence of noise filtering (Gaussian blur, radius of 2 pixels), contrast enhancement (0.2% saturated pixels) and segmentation (Otsu method), after which a watershed algorithm was used to separate closely positioned nuclei. Nuclei positioned at image borders and with abnormal dimensions (due to faulty segmentation) were removed. Ki-67 antibody signal was noise-filtered (Gaussian blur, radius of 2 pixels), background-subtracted (rolling ball radius of 500 pixels) and contrast enhanced up to 0.1% saturated pixels. These images were then used to segment Ki-67 domains (that we interpret as nucleoli) using a fixed threshold of 180 (of 255). Any holes were filled. The segmented domains were used to calculate distance masks from and into Ki-67 domains. For every nucleus, distance maps were overlayed with blurred and background subtracted signals for ^m6^A-Tracer and Ki-67 antibody signals, and per distance group a log2-ratio was calculated over the average nuclear intensity. Finally, nuclei were filtered to have reasonable enrichment scores, which removes cells with faulty segmentation and mitotic cells. Average signal traces were shown for individual nuclei, and averaged between all nuclei of a particular experimental condition.

To calculate Ki-67 depletion efficiency in HCT116 Ki-67-AID cells for the two additional Ki-67 antibodies (Novus and HPA), individual nuclei were segmented as described above. Ki-67 antibody and mClover signals were noise-filtered (Gaussian blur, radius of 2 pixels), median-smoothed (radius of 3 pixels) and background subtracted (rolling window of 250 pixels). Average signals were calculated for every nucleus. For the initial Ki-67 antibody (Abcam), 3D stacks were imaged rather than sections through the middle plane. Image analysis was performed as for the 2D slices, but using 3D implementations instead. 3D Gaussian blurring was performed with a radius of 1 pixel in every direction.

To calculate Ki-67 enrichment at centromeres, individual nuclei were segmented as described above. Centromeres were segmented using a sequence of contrast enhancement (0.1% saturated pixels) and a fixed signal threshold of 100 (of 255). The segmented centromeres were used to calculate distance masks from a centromere. Ki-67 antibody signal was noise-filtered (Gaussian blur, radius of 2 pixels) and background-subtracted (rolling ball radius of 500 pixels). For every nucleus, the distance maps from centromeres were overlayed with blurred and background subtracted Ki-67 signals, and per distance group a log2-ratio was calculated over the average nuclear intensity.

#### Centromere definition

Locations of centromeres were downloaded from the UCSC table browser (group: mapping and sequencing; track: centromeres).

#### External data

**Supplementary Table 1** lists the external data sets that have been used in this study. Previously generated pA-DamID reads were used from the 4DN data repository (https://data.4dnucleome.org/) and processed as described above. These data sets are included in the processed data files that can be found in the GEO repository linked below. Processed Repli-seq data in 5 kb bins were downloaded from the 4DN data repository and the average scores were calculated for 50 kb bins as described above. RNA-seq reads were downloaded from multiple references and used to calculate average FPKM values as described above.

## Data availability

Data sets, computational code and lab journal entries that have been produced for this study are available in the following repositories:

- Data generated in this manuscript: GEO data repository (GSE186206)
- Lab journal records: Open Science Framework (https://osf.io/uqbcr/)

## Code availability

Custom scripts are available on github (https://github.com/vansteensellab/Ki67_dynamics).

## ACKNOWLEDGEMENTS

We thank Masatoshi Takagi for the HCT116 Ki-67-AID cell line, and the NKI Digital Microscopy, Flow Cytometry, Genomics, Protein Production, and RHPC core facilities for technical assistance. We thank members of the BvS lab; Andrew Belmont and other members of the 4D Nucleome Center for Nuclear Cytomics for helpful suggestions. Supported by NIH Common Fund “4D Nucleome” Program grant U54 DK107965 (BvS, DMG), ERC Advanced Grant 694466 (BvS), MSCA/AIRC iCARE2.0 fellowship 800924 (SGM), and Marie Curie Fellowship 838555 (SGM). The Oncode Institute is partly supported by KWF Dutch Cancer Society.

## AUTHOR CONTRIBUTIONS

TvS: Conceived and designed study, conducted majority of experiments and data analysis, wrote manuscript. SGM: Performed experiments. AV: Performed Repli-seq experiments. NQL: Performed ChIP-seq experiments, analyzed data. HT: Performed Hi-C experiments. EdW: Supervised project, analyzed data. DMG: Supervised project. BvS: Designed study, wrote manuscript, supervised project.

## SUPPLEMENTARY FIGURE LEGENDS

**Fig S1.**
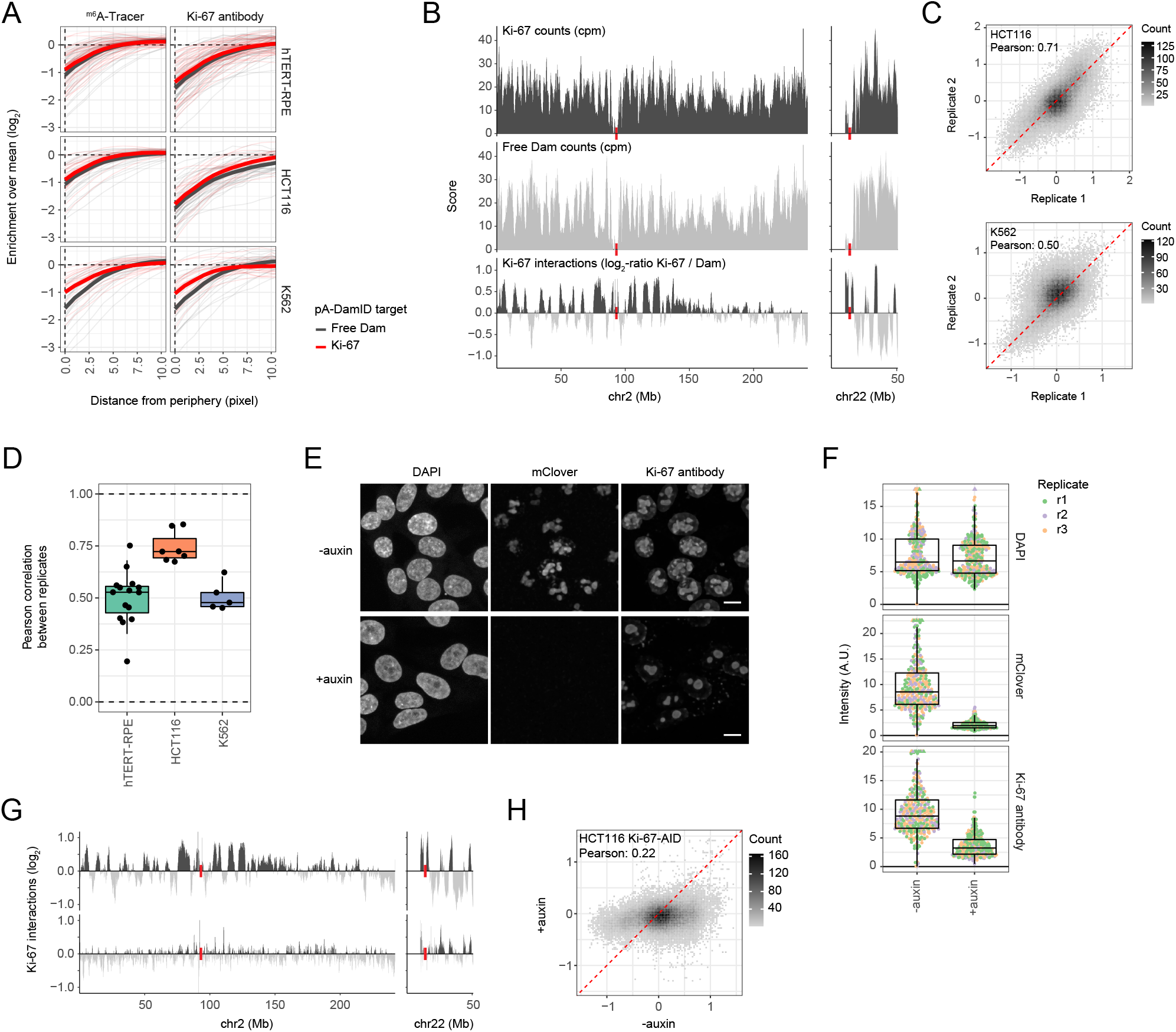
Additional information and controls for Ki-67 pA-DamID. **(A)** Quantification of Ki-67 and ^m6^A-Tracer signals as described in (**Fig 1C**), but instead showing the intensities from the nuclear periphery as defined by DAPI segmentation. **(B)** Example of raw Ki-67 pA-DamID profiles for a representative experiment in HCT116 cells. ^m6^A-marked DNA is specifically amplified, sequenced, counted in 50 kb bins and counts-per-million normalized (cpm) for Ki-67 (top profile) and free Dam (middle profile). A log2-ratio between Ki-67 and Dam accounts for DNA accessibility and amplification bias, and is used as a measure of Ki-67 interactions. Data profiles are smoothed with a running mean across nine 50kb bins for visualization purposes. Centromeres are highlighted by red bars. **(C)** Replicate experiment correlations for Ki-67 interactions (log_2_-ratios) measured in unsynchronized HCT116 cells (top panel) and K562 cells (bottom panel). Pearson correlation scores were calculated with the ‘cor’ function in R. For both examples, p-values are <2.2e-16 as calculated by the ‘cor.test’ function in R. **(D)** Overview of Pearson correlations between replicate experiments for all experimental conditions studied throughout this manuscript, separated by cell line and including perturbation experiments. **(E)** Representative maximum projections of confocal microscopy stacks of HCT116 Ki-67-miniAID-mClover cells (HCT116 Ki-67-AID in short) with normal (top panels) and auxin-mediated depleted Ki-67 levels (bottom panels). Ki-67 levels were visualized directly by mClover fluorescence, and indirectly by immunofluorescence with a Ki-67 antibody. Scale bar: 10 μm. **(F)** Quantification of Ki-67 levels measured by mClover (middle panel) and immunostaining (bottom panel) from three independent replicates. Every point represents a single cell. **(G)** Two representative chromosomes showing the effect of auxin-mediated Ki-67 depletion in HCT116 Ki-67-AID cells on Ki-67 interactions as measured by pA-DamID. Data are averages of two biological replicates and smoothed with a running mean across nine 50kb bins. Centromeres are marked by red bars. **(H)** Correlation of all 50 kb genomic bins from profiles described in (F). As a result of the Ki-67 depletion the standard deviation of the profiled Ki-67 interactions was reduced from 0.40 to 0.19, resulting in a p-value <2.2e-16 (F-test).

**Fig S2.**
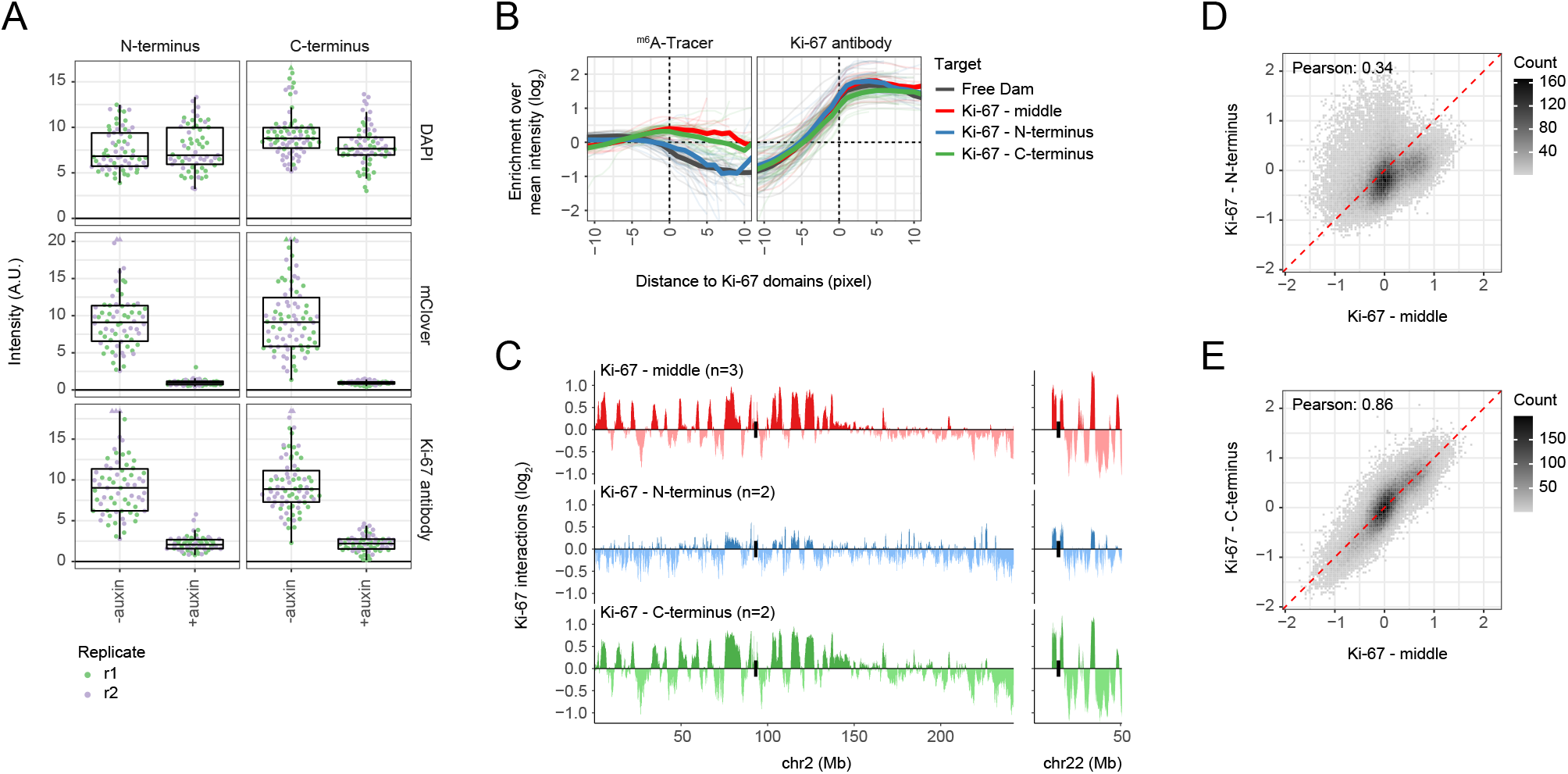
Ki-67 interactions are reproducible between antibodies. **(A)** Quantification of Ki-67 levels as described in (**Fig S1E-F**) based on mClover and Ki-67 immunostaining signals in HCT116 Ki-67-AID cells using different Ki-67 antibodies. For this experiment, single confocal microscopy sections from the middle of cell nuclei were used rather than maximum projections from entire nuclei. Results are combined from two biological replicates. **(B)** Quantification of ^m6^A-Tracer enrichment around Ki-67-marked nucleoli in HCT116 cells, as described in (**Fig 1B-C**). Results are combined from one (middle-targeting antibody) or two (N- and C-terminus antibodies) biological replicates. Data from **Fig 1C** are not included here. **(C)** Ki-67 interactions in HCT116 cells on two representative chromosomes as determined by pA-DamID using three different Ki-67 antibodies. Data profiles are averages of *n* biological replicates and smoothed with a running mean across nine 50kb bins. Centromeres are highlighted by black bars. **(D-E)**Correlation of all 50 kb genomic bins between Ki-67 interactions profiled with the middle-targeting antibody and the N- **(D)** and C-terminus antibodies **(E)**.

**Fig S3.**
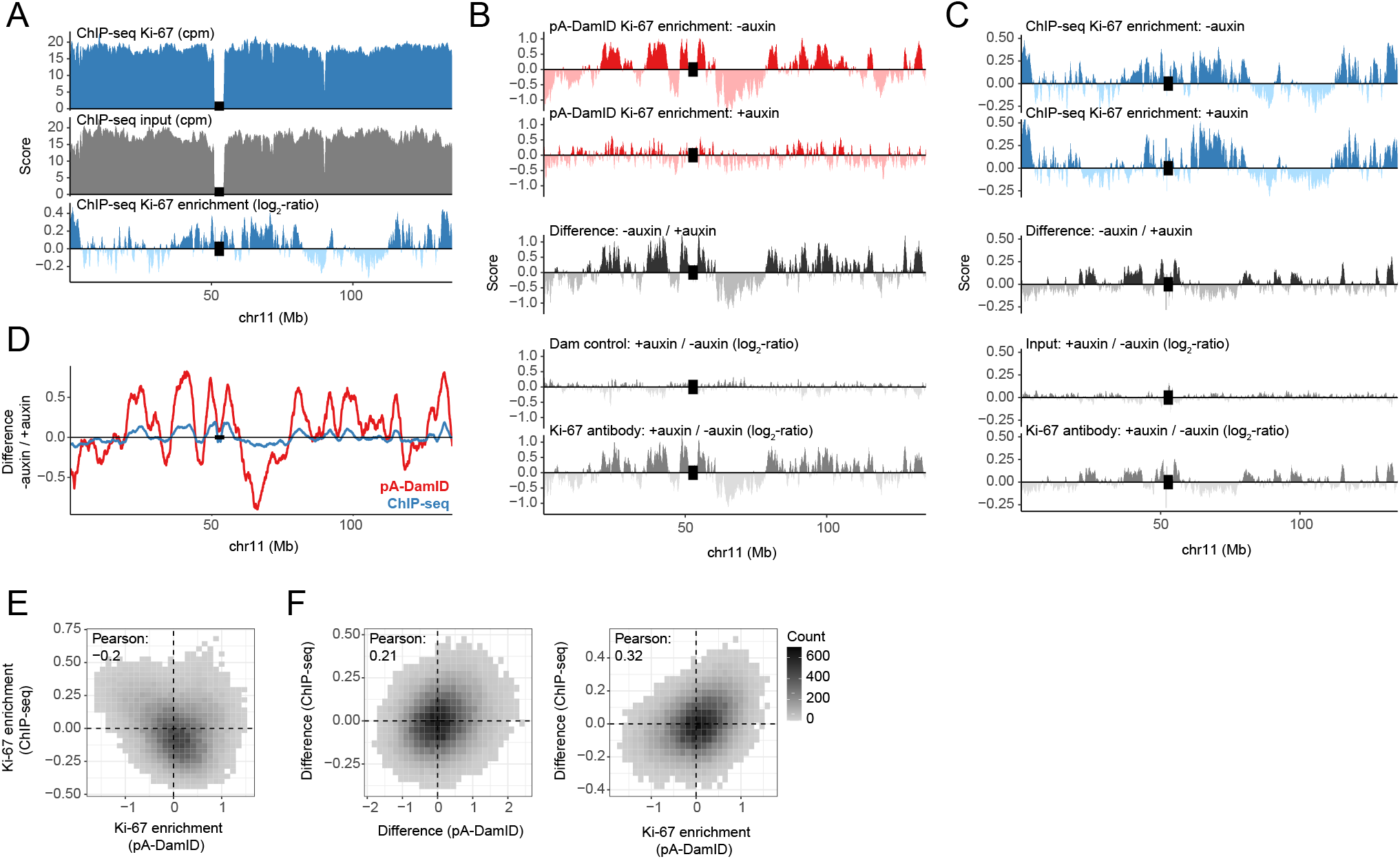
Profiling of Ki-67 interactions with ChIP-seq. **(A)** Example of input and Ki-67 ChIP-seq profiles in HCT116 Ki-67-AID cells and a log_2_-ratio of Ki-67 over input to account for DNA accessibility and amplification bias. Data profiles are smoothed with a running mean across nine 50kb bins for visualization purposes. Centromeres are highlighted by red bars. **(B-C)**Comparison between normalized Ki-67 interactions profiled by pA-DamID (B) and ChIP-seq **(C)**. Data was generated in HCT116 Ki-67-AID cells with and without addition of auxin to deplete Ki-67 *(top panels)*. The difference between these conditions was used to remove any antibody effects or other biases and capture the real signal by Ki-67 *(middle panels)*. Log2-ratios were also calculated between undepleted and depleted conditions using only Dam / input signals or only Ki-67 signals *(bottom panels)*. **(D)** Overlay of the two difference tracks between undepleted and depleted conditions from panels (B-C). Despite a reduced dynamic range, there is a strong concordance between the pA-DamID and ChIP-seq data. **(E)** Binned scatter plot and Pearson correlation between Ki-67 enrichment in undepleted conditions calculated by pA-DamID and ChIP-seq. **(F)** Similar plots as in panel (E), but instead showing the difference between undepleted and depleted conditions *(left panel)* and a comparison of the pA-DamID score in the undepleted condition with the difference as determined by ChIP-seq *(right panel)*.

**Fig S4.**
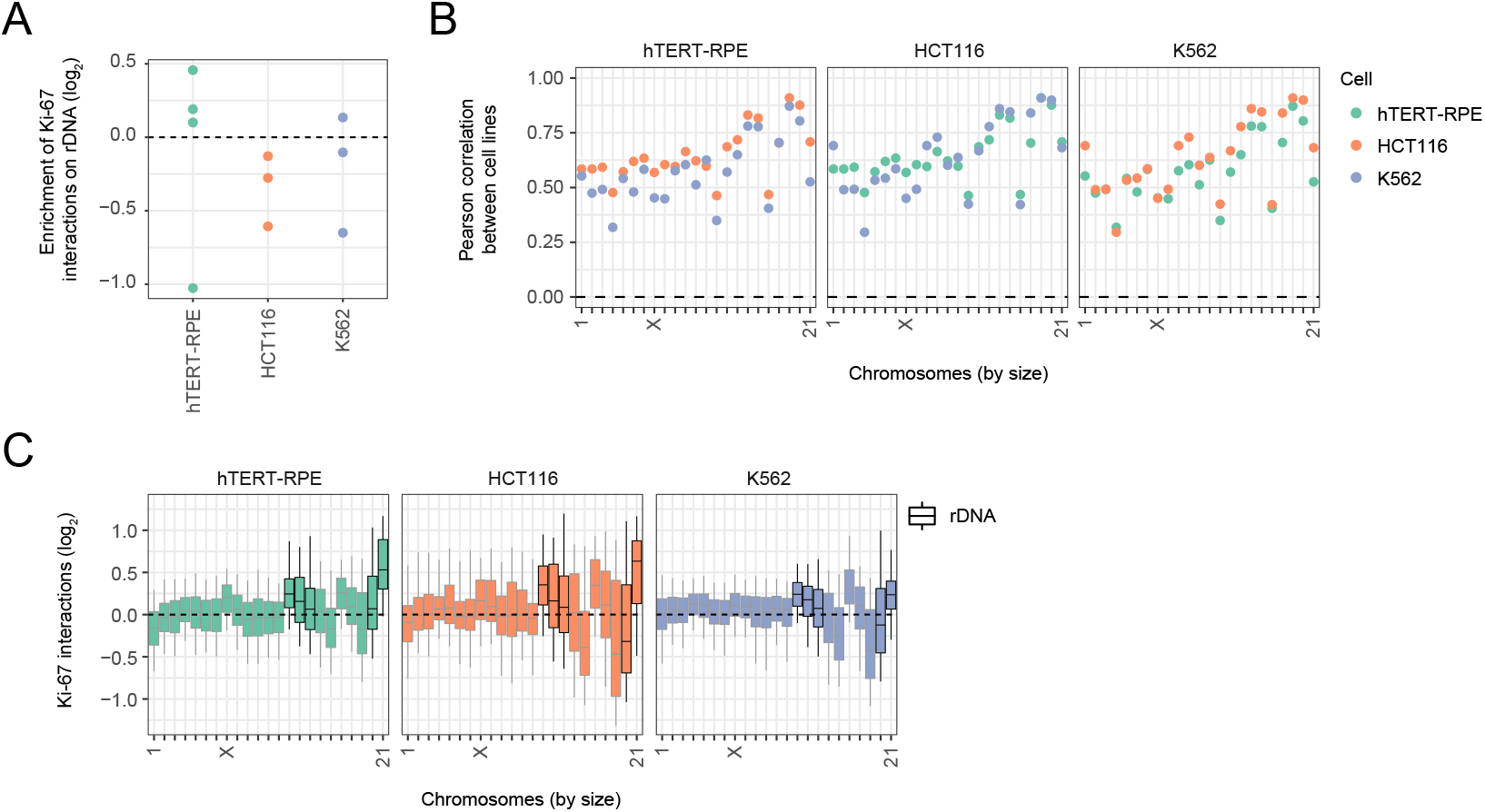
Ki-67 interactions are enriched on small chromosomes, but not enriched at rDNA repeats. **(A)** Enrichment of rDNA sequences detected by Ki-67 pA-DamID compared to the Dam control. Every point represents a biological replicate. **(B)** Pearson correlations between cell lines along all 50 kb genomic bins of individual chromosomes. The panel title indicates the first cell line and the color indicates the second cell line between which the correlation was calculated. For all six cell line comparisons, all Wilcoxon tests between chromosome sizes and the Pearson correlations between cell lines were significant (p-value < 2.4e-13). **(C)** Distributions of Ki-67 interactions for all chromosomes, ordered by decreasing chromosome size. rDNA-containing chromosomes are highlighted by black borders. Boxplots: horizontal lines represent 25^th^, 50^th^, and 75^th^ percentiles; whiskers extend to 5^th^ and 95^th^ percentiles. A Wilcoxon test between chromosome sizes and standard deviations of the Ki-67 interactions was significant (p-value < 2.4e-13) for all three cell lines.

**Fig S5.**
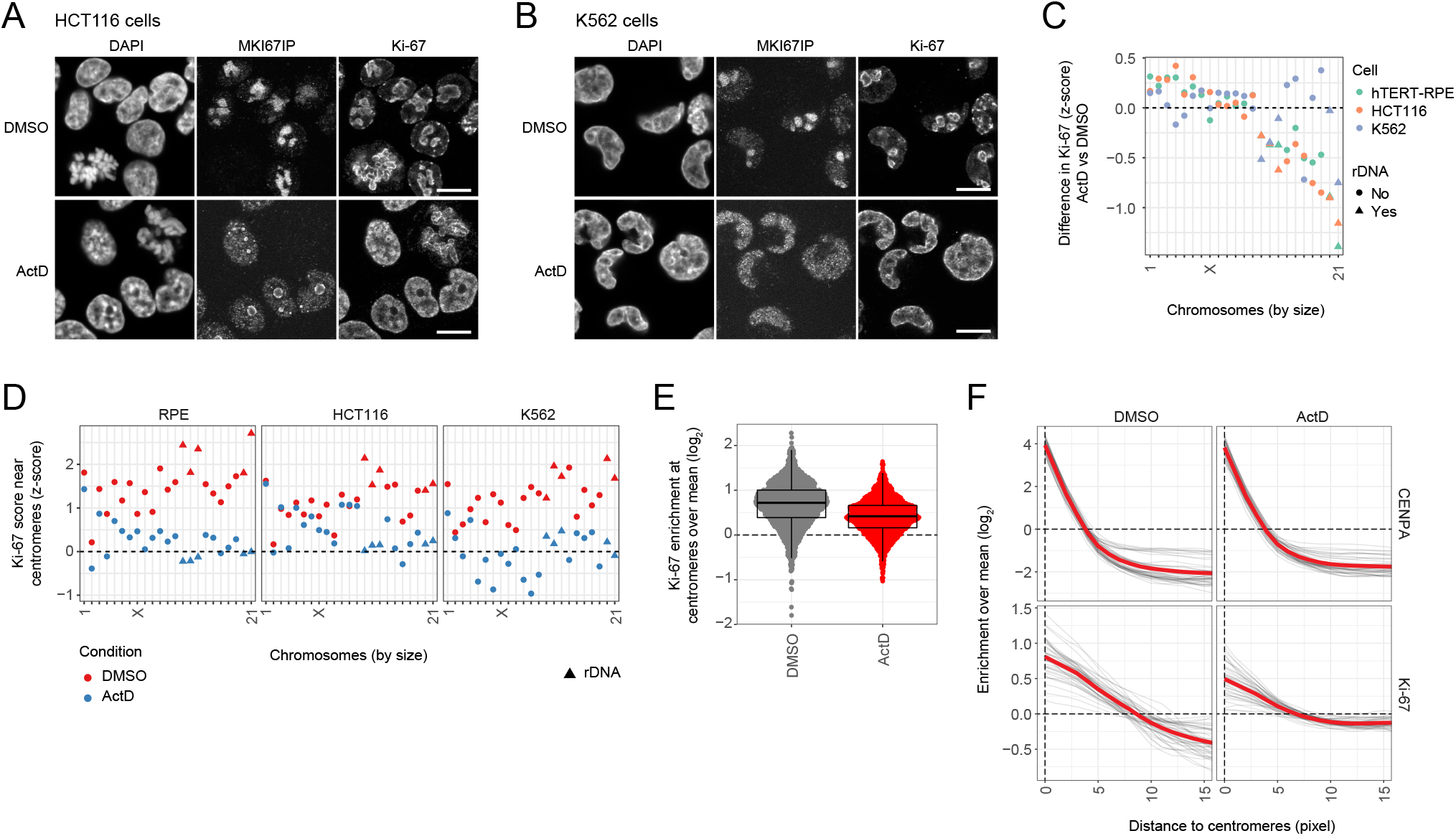
Actinomycin D reduces Ki-67 binding to centromeres. **(A-B)**Representative confocal microscopy sections showing the effect of ActD on nucleolar morphology in HCT116 **(A)** and K562 cells **(B)**, as described in (**Fig 2A**). Scale bar: 10 μm. **(C)** The differences between ActD and control conditions in mean chromosomal Ki-67 signal are plotted for chromosomes sorted by size. rDNA-containing chromosomes are highlighted with triangles. **(D)** The mean Ki-67 interactions scores near centromeres are plotted for each chromosome (within 2Mb of centromeres, overlapping the enrichment in (**Fig 1F**)) in control and ActD conditions. rDNA-containing chromosomes are highlighted in red. **(E-F)**Similar plots showing Ki-67 enrichment at centromeres as (**Fig 1H-I**), but including cells treated with ActD (43 cells in total). DMSO samples are the same as before.

**Fig S6.**
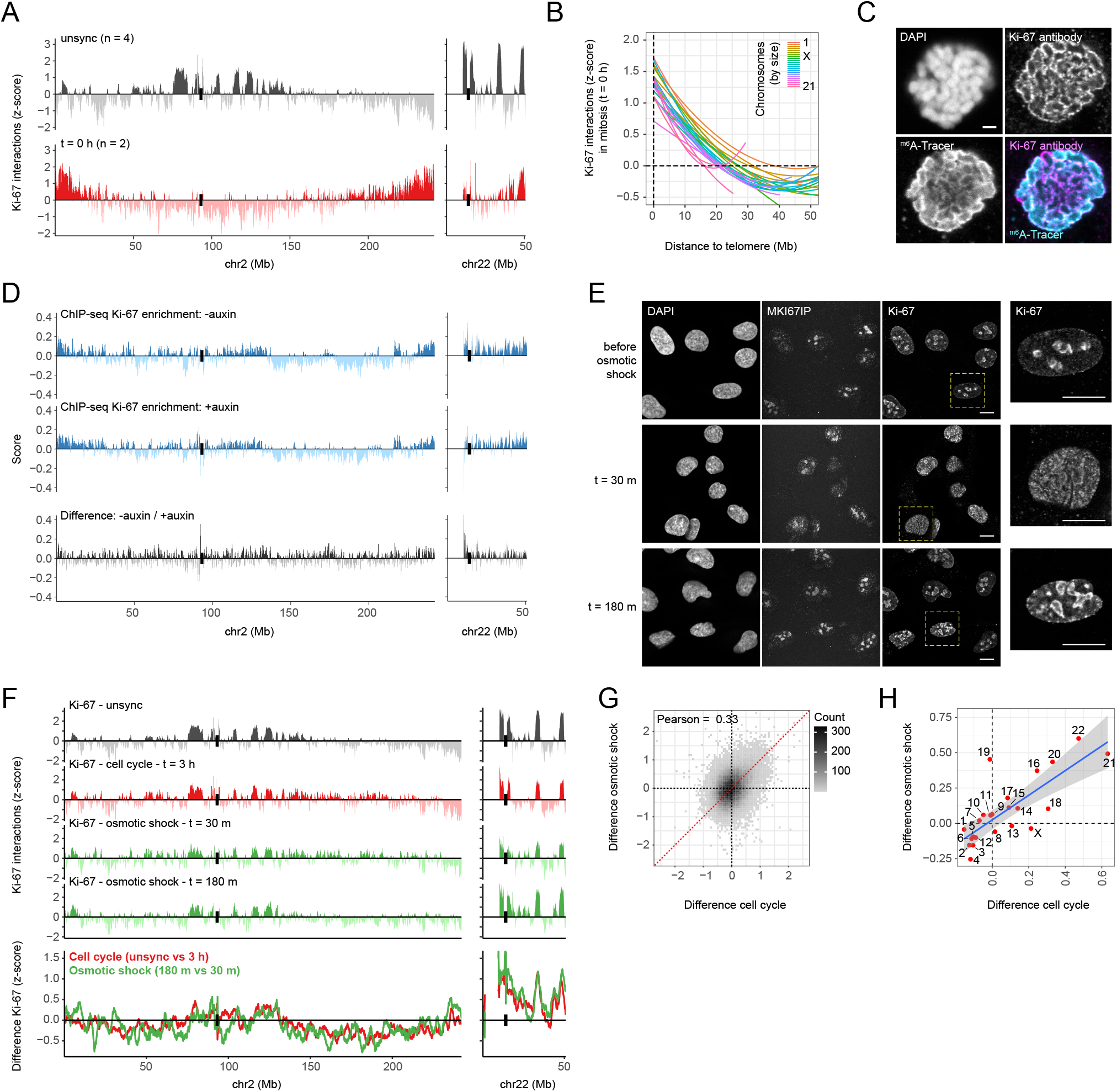
A short osmotic shock recapitulates Ki-67 maturation from PNBs to nucleoli. **(A)** Ki-67 interactions profiles for unsynchronized and metaphase-synchronized (t = 0 h) hTERT-RPE cells. Data profiles are averages of *n* experiments and smoothed with a running mean across nine 50 kb bins. Centromeres are highlighted by black bars. **(B)** Smooth loess curves of Ki-67 interactions plotted against the distance to chromosome ends for cells synchronized in metaphase (t = 0 h). Lines represent individual chromosomes. P-values calculated by Wilcoxon tests between distances to telomeres and Ki-67 interactions are significant for all chromosomes (all <2.2e-16). **(C)** Confocal section of a metaphase hTERT-RPE cell processed with pA-DamID against Ki-67, followed by cytospin centrifugation to create a chromosomal spread and staining against Ki-67 and ^m6^A methylation (see Methods). The middle of the chromosomal territory is not completely stained by the ^m6^A-Tracer, which suggests an incomplete deposition of ^m6^A marks nearby Ki-67 in mitotic cells. Similar depletions in the nuclear interior were never observed for interphase cells processed with Ki-67 pA-DamID. **(D)** Similar data tracks as in (**Fig S3C**), but for cells synchronized in metaphase (t = 0 h). In contract to the pA-DamID data, these data do not support an enrichment of Ki-67 at the distal ends of the chromosomes. **(E)** Representative confocal microscopy sections of hTERT-RPE cells before a 15-minute osmotic shock (0.2x PBS) (top panels), and after 30 and 180 minutes of recovery in complete medium (middle and bottom panels). Formaldehyde-fixed cells were stained for MKI67IP and Ki-67. Scale bar: 10 μm. **(F)** Representative chromosomes showing Ki-67 interactions in unsynchronized hTERT-RPE cells, an early interphase time point (data are the same as in **Fig 2C**) and the two time points following osmotic shock. The difference was calculated between the osmotic shock time points and between early G1 and unsynchronized cells, both of which should resemble the difference between unsynchronized and PNB-enriched cells. Data profiles are averages of two experiments and smoothed with a running mean across nine 50 kb bins. Centromeres are highlighted by black bars. **(G)** Scatter plot and Pearson correlation between the two difference tracks from panel (F). The ‘cor.test’ function in R was used to determine statistical significance, resulting in a p-value <2.2e-16. **(H)** Changes in chromosomal distributions of Ki-67 are similar for progression in G1 phase (x-axis: difference in mean interactions per chromosome between the unsynchronized cells and the 3 h time point) and upon recovery from osmotic shock (y-axis: difference in mean interactions per chromosome between 180 min and 30 min after termination of the osmotic shock (OS)). The blue line represents a linear model with a 95%-confidence interval.

**Fig S7.**
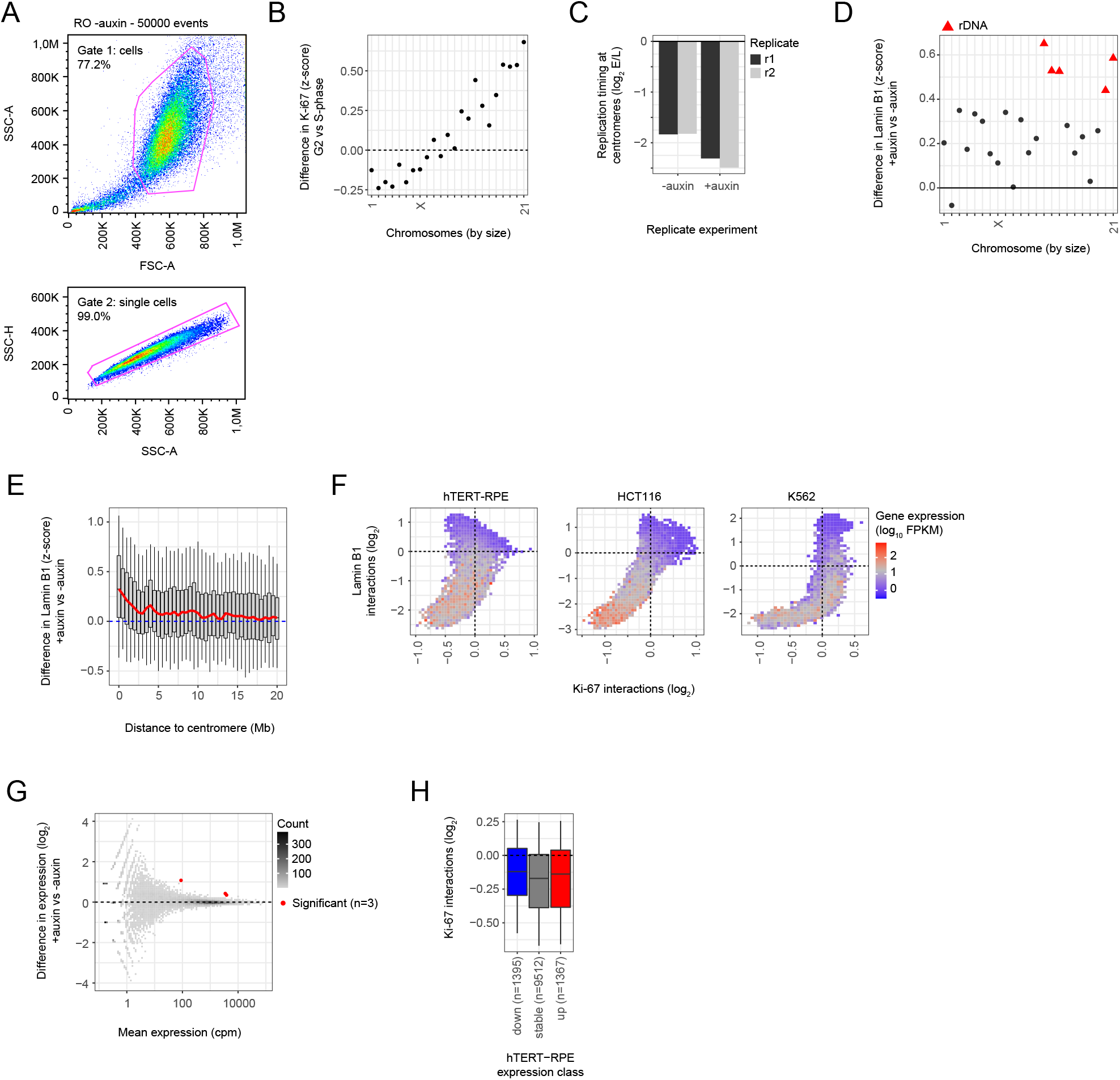
Loss of Ki-67 interactions does not correlate with gene expression changes. **(A)** Gating strategy for FACS analysis of synchronized HCT116 Ki-67-AID cells, showing 50,000 events for G2 control cells (RO -auxin condition). First, cells were selected using a gate based on forward scatter area (FSC-A) and side scatter area (SSC-A). Second, single cells were selected using a gate based on SSC-A and side scatter height (SSC-H). At least 30,000 single cells were selected for all samples. **(B)** Differences in mean Ki-67 chromosome scores between S- and G2-synchronized HCT116 Ki-67-AID cells. **(C)** Bar plot showing average replication timing (log2-ratio Early/Late) for centromeres. All reads were selected that aligned to regions marked as centromeres (UCSC Table browser; hg38 centromeres) regardless of their mapping quality and, following counts-per-million normalization, the E/L ratio was calculated between the total centromeric counts. Between 3-5% of the late-fraction mapped to centromeres. **(D-E)**Similar plots as (**Fig 5F-G**), but showing differences in Lamin B1 interactions instead. **(F)** Similar plot as (**Fig 4C**), but instead highlighting for every gene the expression level (in log_10_ FPKM). Expression data are used from (Consortium, 2012; Dai *et al*, 2018; Durrbaum *et al*, 2018; Harenza *et al*, 2017; Kelso *et al*, 2017; Slaats *et al*, 2015; Sun *et al.*, 2017) (see Methods). **(G)** Scatterplot of mean gene expression (counts per million; cpm) versus expression difference (log_2_-fold change) following auxin-mediated Ki-67 depletion in HCT116 Ki-67-AID cells. Significant genes (Benjamini-Hochberg adjusted p-value < 0.05) are highlighted in red. **(H)** Boxplots showing Ki-67 interactions for differentially expressed genes following Ki-67 depletion in hTERT-RPE cells. Gene expression calls are from (Sun *et al.*, 2017). Boxplots: horizontal lines represent 25th, 50th, and 75th percentiles; whiskers extend to 5th and 95th percentiles.

**Fig S8.**
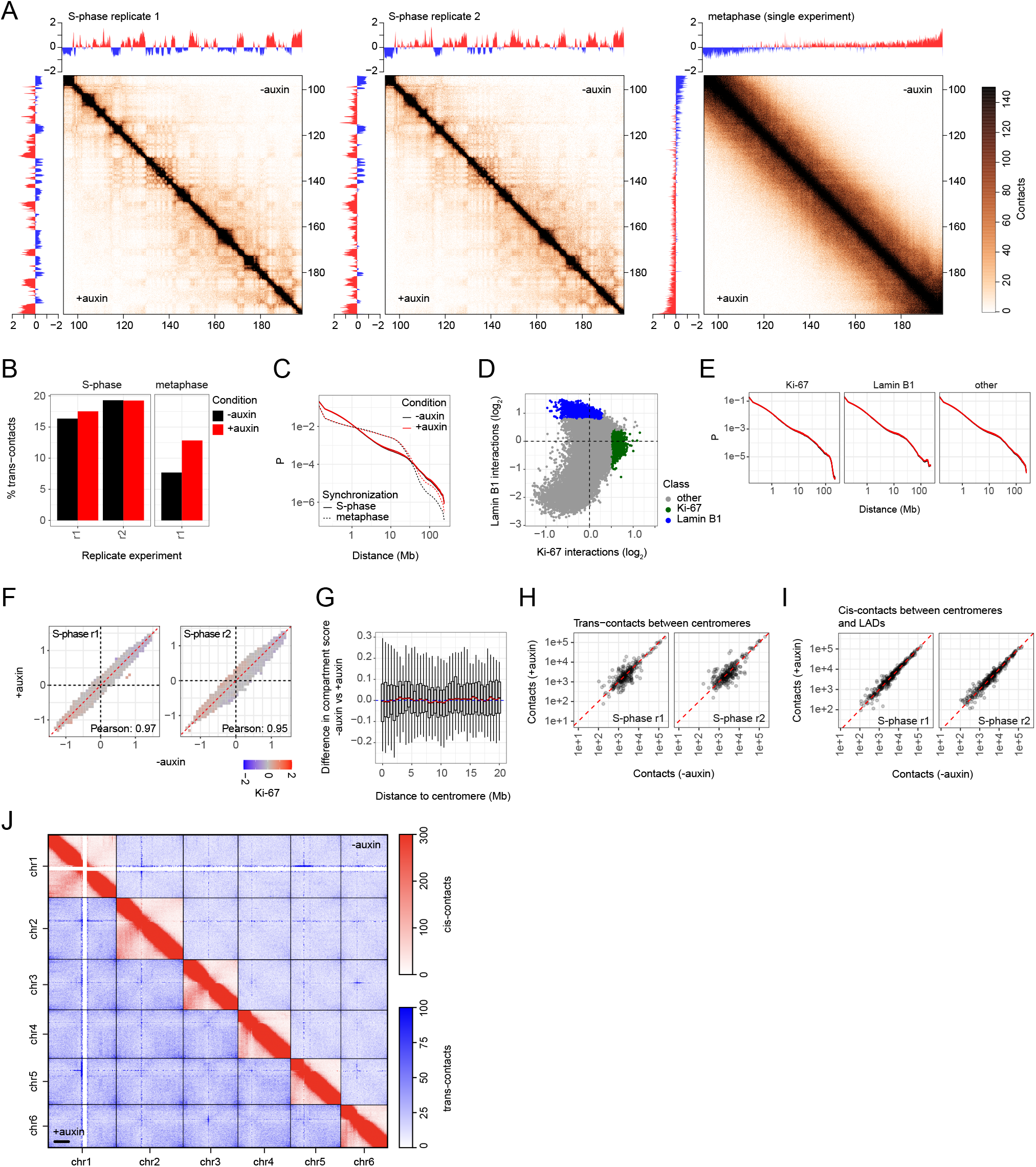
Ki-67 depletion in interphase does not affect genome contacts. **(A)** Representative chromosome arm showing genome contacts captured by Hi-C in HCT116 Ki-67-AID cells with and without addition of auxin to deplete Ki-67 (resolution: 100 kb bins). Two replicates were generated in S-phase synchronized cells *(left and middle panels)* and one replicate for metaphase cells *(right panel)*. Compartment scores calculated from the data are included on the sides. **(B)** Fraction of trans-contacts for the samples described in panel (A). **(C)** Genome-wide curves of contact probability (P) versus genomic distance. **(D-E)** Genomic regions positioned in the two ends of the hammer-head were selected to select domains preferentially at the nuclear lamina and near Ki-67 **(D)**, and used to determine genome-wide contact probability curves within these regions **(E)**. Loss of Ki-67 binding does not affect genome contacts at the domains that were initially bound by Ki-67. **(F)** Similar plot as (**Fig 5C**), but instead showing a binned scatter plot with compartment scores on the axes. Genomic regions bound by Ki-67 do not change their compartment score upon auxin-mediated depletion. **(G)** Similar plot as (**Fig 5F**), but instead showing the difference in compartment score upon depletion of Ki-67. Loss of Ki-67 binding and decreased replication timing does not affect the compartment score. **(H)** Trans-contacts between genomic domains flanking the centromere (within 2 Mb) were calculated, for undepleted and depleted conditions. Every point represents a chromosome pair. Loss of Ki-67 binding and nucleolar positioning for centromeres (Sobecki *et al.*, 2016) does not affect centromere clustering as measured by Hi-C. **(I)** Cis-contacts were calculated between genomic domains flanking the centromere (within 2 Mb) and LADs, for undepleted and depleted conditions. Every point represents the combined contacts between a centromere and one LAD. Loss of Ki-67 binding and increased peripheral positioning of centromeres (**Fig S7E**) does not affect long-range DNA contacts at the nuclear lamina. **(J)** Overview of cis- and trans-contacts for six chromosomes from the mitotic Hi-C samples, showing the contacts in normal *(top right)* and depleted conditions *(bottom left)* (resolution: 1 Mb bins). Loss of Ki-67 in mitosis increases trans-contacts and contacts perpendicular to the centromere, which could correspond to increased folding of the chromosome arms on each other.

## REFERENCES

Barra V, Fachinetti D (2018) The dark side of centromeres: types, causes and consequences of structural abnormalities implicating centromeric DNA. Nat Commun 9: 4340

Bizhanova A, Kaufman PD (2021) Close to the edge: Heterochromatin at the nucleolar and nuclear peripheries. Biochim Biophys Acta Gene Regul Mech 1864: 194666

Bloom K, Costanzo V (2017) Centromere Structure and Function. Prog Mol Subcell Biol 56: 515–539

Bolzer A, Kreth G, Solovei I, Koehler D, Saracoglu K, Fauth C, Muller S, Eils R, Cremer C, Speicher MR et al (2005) Three-dimensional maps of all chromosomes in human male fibroblast nuclei and prometaphase rosettes. PLoS Biol 3: e157

Booth DG, Takagi M, Sanchez-Pulido L, Petfalski E, Vargiu G, Samejima K, Imamoto N, Ponting CP, Tollervey D, Earnshaw WC et al (2014) Ki-67 is a PP1-interacting protein that organises the mitotic chromosome periphery. Elife 3: e01641

Bridger JM, Kill IR, Lichter P (1998) Association of pKi-67 with satellite DNA of the human genome in early G1 cells. Chromosome Res 6: 13–24

Bury L, Moodie B, Ly J, McKay LS, Miga KH, Cheeseman IM (2020) Alpha-satellite RNA transcripts are repressed by centromere-nucleolus associations. Elife 9

Carron C, Balor S, Delavoie F, Plisson-Chastang C, Faubladier M, Gleizes PE, O’Donohue MF (2012) Post-mitotic dynamics of pre-nucleolar bodies is driven by pre-rRNA processing. J Cell Sci 125: 4532–4542

Chen S, Zhou Y, Chen Y, Gu J (2018) fastp: an ultra-fast all-in-one FASTQ preprocessor. Bioinformatics 34: i884–i890

Consortium EP (2012) An integrated encyclopedia of DNA elements in the human genome. Nature 489: 57–74

Cuylen S, Blaukopf C, Politi AZ, Muller-Reichert T, Neumann B, Poser I, Ellenberg J, Hyman AA, Gerlich DW (2016) Ki-67 acts as a biological surfactant to disperse mitotic chromosomes. Nature 535: 308–312

Cuylen-Haering S, Petrovic M, Hernandez-Armendariz A, Schneider MWG, Samwer M, Blaukopf C, Holt LJ, Gerlich DW (2020) Chromosome clustering by Ki-67 excludes cytoplasm during nuclear assembly. Nature 587: 285–290

Dai Z, Mentch SJ, Gao X, Nichenametla SN, Locasale JW (2018) Methionine metabolism influences genomic architecture and gene expression through H3K4me3 peak width. Nat Commun 9: 1955

Dekker J, Belmont AS, Guttman M, Leshyk VO, Lis JT, Lomvardas S, Mirny LA, O’Shea CC, Park PJ, Ren B et al (2017) The 4D nucleome project. Nature 549: 219–226

Dillinger S, Straub T, Nemeth A (2017) Nucleolus association of chromosomal domains is largely maintained in cellular senescence despite massive nuclear reorganisation. PLoS One 12: e0178821

Dobin A, Davis CA, Schlesinger F, Drenkow J, Zaleski C, Jha S, Batut P, Chaisson M, Gingeras TR (2013) STAR: ultrafast universal RNA-seq aligner. Bioinformatics 29: 15–21

Dundr M, Misteli T, Olson MO (2000) The dynamics of postmitotic reassembly of the nucleolus. J Cell Biol 150: 433–446

Durrbaum M, Kruse C, Nieken KJ, Habermann B, Storchova Z (2018) The deregulated microRNAome contributes to the cellular response to aneuploidy. BMC Genomics 19: 197

Elbatsh AMO, Kim E, Eeftens JM, Raaijmakers JA, van der Weide RH, Garcia-Nieto A, Bravo S, Ganji M, Uit de Bos J, Teunissen H et al (2019) Distinct Roles for Condensin’s Two ATPase Sites in Chromosome Condensation. Mol Cell 76: 724–737 e725

Falk M, Feodorova Y, Naumova N, Imakaev M, Lajoie BR, Leonhardt H, Joffe B, Dekker J, Fudenberg G, Solovei I et al (2019) Heterochromatin drives compartmentalization of inverted and conventional nuclei. Nature 570: 395–399

Garwain O, Sun X, Iyer DR, Li R, Zhu LJ, Kaufman PD (2020) The chromatin-binding domain of Ki-67 together with p53 protects human chromosomes from mitotic damage. bioRxiv: 2020.2010.2016.342352

Gerdes J, Lemke H, Baisch H, Wacker HH, Schwab U, Stein H (1984) Cell cycle analysis of a cell proliferation-associated human nuclear antigen defined by the monoclonal antibody Ki-67. J Immunol 133: 1710–1715

Giunta S, Herve S, White RR, Wilhelm T, Dumont M, Scelfo A, Gamba R, Wong CK, Rancati G, Smogorzewska A et al (2021) CENP-A chromatin prevents replication stress at centromeres to avoid structural aneuploidy. Proc Natl Acad Sci U S A 118

Greil F, Moorman C, van Steensel B (2006) DamID: mapping of in vivo protein-genome interactions using tethered DNA adenine methyltransferase. Methods Enzymol 410: 342–359

Guelen L, Pagie L, Brasset E, Meuleman W, Faza MB, Talhout W, Eussen BH, de Klein A, Wessels L, de Laat W et al (2008) Domain organization of human chromosomes revealed by mapping of nuclear lamina interactions. Nature 453: 948–951

Haarhuis JHI, van der Weide RH, Blomen VA, Yanez-Cuna JO, Amendola M, van Ruiten MS, Krijger PHL, Teunissen H, Medema RH, van Steensel B et al (2017) The Cohesin Release Factor WAPL Restricts Chromatin Loop Extension. Cell 169: 693–707 e614

Harenza JL, Diamond MA, Adams RN, Song MM, Davidson HL, Hart LS, Dent MH, Fortina P, Reynolds CP, Maris JM (2017) Transcriptomic profiling of 39 commonly-used neuroblastoma cell lines. Sci Data 4: 170033

Hiratani I, Ryba T, Itoh M, Yokochi T, Schwaiger M, Chang CW, Lyou Y, Townes TM, Schubeler D, Gilbert DM (2008) Global reorganization of replication domains during embryonic stem cell differentiation. PLoS Biol 6: e245

Kelso TWR, Porter DK, Amaral ML, Shokhirev MN, Benner C, Hargreaves DC (2017) Chromatin accessibility underlies synthetic lethality of SWI/SNF subunits in ARID1A-mutant cancers. Elife 6

Kind J, Pagie L, Ortabozkoyun H, Boyle S, de Vries SS, Janssen H, Amendola M, Nolen LD, Bickmore WA, van Steensel B (2013) Single-cell dynamics of genome-nuclear lamina interactions. Cell 153: 178–192

Leemans C, van der Zwalm MCH, Brueckner L, Comoglio F, van Schaik T, Pagie L, van Arensbergen J, van Steensel B (2019) Promoter-Intrinsic and Local Chromatin Features Determine Gene Repression in LADs. Cell 177: 852–864 e814

Lima-De-Faria A, Reitalu J (1963) Heterochromatin in human male leukocytes. J Cell Biol 16: 315–322

Liu NQ, Maresca M, van den Brand T, Braccioli L, Schijns M, Teunissen H, Bruneau BG, Nora EP, de Wit E (2021) WAPL maintains a cohesin loading cycle to preserve cell-type-specific distal gene regulation. Nat Genet 53: 100–109

Love MI, Huber W, Anders S (2014) Moderated estimation of fold change and dispersion for RNA-seq data with DESeq2. Genome Biol 15: 550

Luderus ME, den Blaauwen JL, de Smit OJ, Compton DA, van Driel R (1994) Binding of matrix attachment regions to lamin polymers involves single-stranded regions and the minor groove. Mol Cell Biol 14: 6297–6305

MacCallum DE, Hall PA (1999) Biochemical characterization of pKi67 with the identification of a mitotic-specific form associated with hyperphosphorylation and altered DNA binding. Exp Cell Res 252: 186–198

MacCallum DE, Hall PA (2000) The biochemical characterization of the DNA binding activity of pKi67. J Pathol 191: 286–298

Marchal C, Sasaki T, Vera D, Wilson K, Sima J, Rivera-Mulia JC, Trevilla-Garcia C, Nogues C, Nafie E, Gilbert DM (2018) Genome-wide analysis of replication timing by next-generation sequencing with E/L Repli-seq. Nat Protoc 13: 819–839

Marchal C, Sima J, Gilbert DM (2019) Control of DNA replication timing in the 3D genome. Nat Rev Mol Cell Biol 20: 721–737

Matheson TD, Kaufman PD (2017) The p150N domain of chromatin assembly factor-1 regulates Ki-67 accumulation on the mitotic perichromosomal layer. Mol Biol Cell 28: 21–29

Meuleman W, Peric-Hupkes D, Kind J, Beaudry JB, Pagie L, Kellis M, Reinders M, Wessels L, van Steensel B (2013) Constitutive nuclear lamina-genome interactions are highly conserved and associated with A/T-rich sequence. Genome Res 23: 270–280

Miller I, Min M, Yang C, Tian C, Gookin S, Carter D, Spencer SL (2018) Ki67 is a Graded Rather than a Binary Marker of Proliferation versus Quiescence. Cell Rep 24: 1105–1112 e1105

Mrouj K, Andres-Sanchez N, Dubra G, Singh P, Sobecki M, Chahar D, Al Ghoul E, Aznar AB, Prieto S, Pirot N et al (2021) Ki-67 regulates global gene expression and promotes sequential stages of carcinogenesis. Proc Natl Acad Sci U S A 118

Nemeth A, Grummt I (2018) Dynamic regulation of nucleolar architecture. Curr Opin Cell Biol 52: 105–111

Ochs RL, Lischwe MA, Shen E, Carroll RE, Busch H (1985) Nucleologenesis: composition and fate of prenucleolar bodies. Chromosoma 92: 330–336

Ohno S, Kaplan WD, Kinosita R (1959) The centromeric and nucleolus-associated heterochromatin of Rattus norvegicus. Exp Cell Res 16: 348–357

Perry RP, Kelley DE (1970) Inhibition of RNA synthesis by actinomycin D: characteristic dose-response of different RNA species. J Cell Physiol 76: 127–139

Politz JCR, Scalzo D, Groudine M (2016) The redundancy of the mammalian heterochromatic compartment. Curr Opin Genet Dev 37: 1–8

Quinodoz SA, Ollikainen N, Tabak B, Palla A, Schmidt JM, Detmar E, Lai MM, Shishkin AA, Bhat P, Takei Y et al (2018) Higher-Order Inter-chromosomal Hubs Shape 3D Genome Organization in the Nucleus. Cell 174: 744–757 e724

Ragoczy T, Telling A, Scalzo D, Kooperberg C, Groudine M (2014) Functional redundancy in the nuclear compartmentalization of the late-replicating genome. Nucleus 5: 626–635

Reddy KL, Zullo JM, Bertolino E, Singh H (2008) Transcriptional repression mediated by repositioning of genes to the nuclear lamina. Nature 452: 243–247

Remnant L, Kochanova NY, Reid C, Cisneros-Soberanis F, Earnshaw WC (2021) The intrinsically disorderly story of Ki-67. Open Biol 11: 210120

Saiwaki T, Kotera I, Sasaki M, Takagi M, Yoneda Y (2005) In vivo dynamics and kinetics of pKi-67: transition from a mobile to an immobile form at the onset of anaphase. Exp Cell Res 308: 123–134

Savino TM, Gebrane-Younes J, De Mey J, Sibarita JB, Hernandez-Verdun D (2001) Nucleolar assembly of the rRNA processing machinery in living cells. J Cell Biol 153: 1097–1110

Scholzen T, Gerdes J (2000) The Ki-67 protein: from the known and the unknown. J Cell Physiol 182: 311–322

Servant N, Varoquaux N, Lajoie BR, Viara E, Chen CJ, Vert JP, Heard E, Dekker J, Barillot E (2015) HiC-Pro: an optimized and flexible pipeline for Hi-C data processing. Genome Biol 16: 259

Slaats GG, Wheway G, Foletto V, Szymanska K, van Balkom BW, Logister I, Den Ouden K, Keijzer-Veen MG, Lilien MR, Knoers NV et al (2015) Screen-based identification and validation of four new ion channels as regulators of renal ciliogenesis. J Cell Sci 128: 4550–4559

Sobecki M, Mrouj K, Camasses A, Parisis N, Nicolas E, Lleres D, Gerbe F, Prieto S, Krasinska L, David A et al (2016) The cell proliferation antigen Ki-67 organises heterochromatin. Elife 5: e13722

Sobecki M, Mrouj K, Colinge J, Gerbe F, Jay P, Krasinska L, Dulic V, Fisher D (2017) Cell-Cycle Regulation Accounts for Variability in Ki-67 Expression Levels. Cancer Res 77: 2722–2734

Solovei I, Wang AS, Thanisch K, Schmidt CS, Krebs S, Zwerger M, Cohen TV, Devys D, Foisner R, Peichl L et al (2013) LBR and lamin A/C sequentially tether peripheral heterochromatin and inversely regulate differentiation. Cell 152: 584–598

Stenstrom L, Mahdessian D, Gnann C, Cesnik AJ, Ouyang W, Leonetti MD, Uhlen M, Cuylen-Haering S, Thul PJ, Lundberg E (2020) Mapping the nucleolar proteome reveals a spatiotemporal organization related to intrinsic protein disorder. Mol Syst Biol 16: e9469

Su JH, Zheng P, Kinrot SS, Bintu B, Zhuang X (2020) Genome-Scale Imaging of the 3D Organization and Transcriptional Activity of Chromatin. Cell 182: ‘641–1659 e1626

Sun X, Bizhanova A, Matheson TD, Yu J, Zhu LJ, Kaufman PD (2017) Ki-67 Contributes to Normal Cell Cycle Progression and Inactive X Heterochromatin in p21 Checkpoint-Proficient Human Cells. Mol Cell Biol 37

Sun X, Kaufman PD (2018) Ki-67: more than a proliferation marker. Chromosoma 127: 175–186

Takagi M, Natsume T, Kanemaki MT, Imamoto N (2016) Perichromosomal protein Ki67 supports mitotic chromosome architecture. Genes Cells 21: 1113–1124

van der Weide RH, van den Brand T, Haarhuis JHI, Teunissen H, Rowland BD, de Wit E (2021) Hi-C analyses with GENOVA: a case study with cohesin variants. NAR Genom Bioinform 3: lqab040

van Koningsbruggen S, Gierlinski M, Schofield P, Martin D, Barton GJ, Ariyurek Y, den Dunnen JT, Lamond AI (2010) High-resolution whole-genome sequencing reveals that specific chromatin domains from most human chromosomes associate with nucleoli. Mol Biol Cell 21: 3735–3748

van Schaik T, Vos M, Peric-Hupkes D, Hn Celie P, van Steensel B (2020) Cell cycle dynamics of lamina-associated DNA. EMBO Rep 21: e50636

van Steensel B, Belmont AS (2017) Lamina-Associated Domains: Links with Chromosome Architecture, Heterochromatin, and Gene Repression. Cell 169: 780–791

Verheijen R, Kuijpers HJ, van Driel R, Beck JL, van Dierendonck JH, Brakenhoff GJ, Ramaekers FC (1989) Ki-67 detects a nuclear matrix-associated proliferation-related antigen. II. Localization in mitotic cells and association with chromosomes. J Cell Sci 92 (Pt 4): 531–540

Vertii A, Ou J, Yu J, Yan A, Pages H, Liu H, Zhu LJ, Kaufman PD (2019) Two contrasting classes of nucleolus-associated domains in mouse fibroblast heterochromatin. Genome Res 29: 1235–1249

Vogel MJ, Peric-Hupkes D, van Steensel B (2007) Detection of in vivo protein-DNA interactions using DamID in mammalian cells. Nat Protoc 2: 1467–1478

Yamazaki H, Takagi M, Kosako H, Hirano T, Yoshimura SH (2022) Cell cycle-specific phase separation regulated by protein charge blockiness. Nat Cell Biol 24: 625–632

Zatsepina OV, Dudnic OA, Todorov IT, Thiry M, Spring H, Trendelenburg MF (1997) Experimental induction of prenucleolar bodies (PNBs) in interphase cells: interphase PNBs show similar characteristics as those typically observed at telophase of mitosis in untreated cells. Chromosoma 105: 418–430

Zhang LF, Huynh KD, Lee JT (2007) Perinucleolar targeting of the inactive X during S phase: evidence for a role in the maintenance of silencing. Cell 129: 693–706

