## Supplementary Table 1 for "Dynamic chromosomal interactions and control of heterochromatin positioning by Ki-67"

**External data sets used**

| Data type | Cell type | Target | Reference | PMID | ID |
|---|---|---|---|---|---|
| pA-DamID | hTERT-RPE | LaminB1 | van Schaik, 2020; 4D Nucleome | 32893442 | 4DNESENITEIC |
| pA-DamID | HCT116 | LaminB1 | van Schaik, 2020; 4D Nucleome | 32893442 | 4DNESWB729QB |
| pA-DamID | K562 | LaminB1 | van Schaik, 2020; 4D Nucleome | 32893442 | 4DNESUMP6SS1 |
| pA-DamID | K562 | H3K27me3 | van Schaik, 2020; 4D Nucleome | 32893442 | 4DNES7YUHPFR |
| Repli-seq | hTERT-RPE | | 4D Nucleome; Gilbert lab | 28905911 | 4DNES674QWXK, 4DNESUNOW1OZ |
| Repli-seq | HCT116 | | 4D Nucleome; Gilbert lab | 28905911 | 4DNESLC8TDK4, 4DNESSYKYIK3 |
| Repli-seq | K562 | | 4D Nucleome; Gilbert lab | 28905911 | 4DNES1GSPUT8, 4DNES9YA22WT |
| RNA-seq | hTERT-RPE | | Slaats, 2015; SRA | 26546361 | SRX1411451 |
| RNA-seq | hTERT-RPE | | Sun, 2017; SRA | 28630280 | SRX2805061, SRX2172520 |
| RNA-seq | hTERT-RPE | | Dürrbaum, 2018; GEO | 29703144 | GSM2747191, GSM2747192, GSM2747193 |
| RNA-seq | hTERT-RPE | | Harenza, 2017; GEO | 28350380 | GSM2371252 |
| RNA-seq | HCT116 | | ENCODE | 22955616 | ENCFF000DKT, ENCFF000DKW, ENCFF000DKV, ENCFF000DKX, ENCFF000DKY, ENCFF000DKU |
| RNA-seq | HCT116 | | Kelso, 2017; GEO | 28967863 | GSM2719768, GSM2719769 |
| RNA-seq | HCT116 | | Dai, 2018; GEO | 29769529 | GSM2775145, GSM2775146 |
| RNA-seq | K562 | | ENCODE | 22955616 | ENCFF001RED, ENCFF001REG, ENCFF001RWD, ENCFF001RVV, ENCFF001RWE, ENCFF001RWF, ENCFF001RDD, ENCFF001RDE, ENCFF000HFF, ENCFF000HFH |
